# Capturing the imagination: mapping imagery ability across our multidimensional sense of touch

**DOI:** 10.64898/2026.08.26.747318

**Authors:** Renee Lustenhouwer, H. Chris Dijkerman

**Author notes:** corresponding author Lustenhouwer, R. (Renee). Dijkerman, H.C. (Chris).

## Abstract

Mental imagery relies on modality-specific representations and varies across individuals. The tactile imagery modality remains relatively understudied. Though imagined and actual touch show perceptual and neural overlap, previous work has largely examined specific touch types in isolation. Likewise, imagery ability is often equated with imagery vividness, omitting other imagery processing components. Here we aimed to provide a comprehensive assessment of tactile imagery in terms of touch types – object properties for active touch, different tactile sensations across body sites for passive touch – and imagery processing components. 136 healthy adults imagined various touch types, and rated imagery vividness and their ability to maintain and transform each sensation on 5-point Likert scales. A subsample (N=46) additionally rated pleasantness on Visual Analogue Scales. Overall, most participants reported at least some imagery, though individual variability was substantial. Imagery ability varied significantly across objects and properties, with specific object-property pairs eliciting stronger imagery than others. Imagery was significantly stronger for body sites with higher receptor density (i.e. lip and foot), and imagined stroking was significantly weaker than vibration and pinching. Pinching was significantly less pleasant than stroking and vibration. Pleasantness and imagery ability showed significant positive correlations for active touch and passive stroking. Active and passive imagery ability were strongly positively correlated, as were the imagery processing components. These findings demonstrate that tactile imagery is not a unitary ability. Instead, it varies across individuals and tactile sensations, mirroring the multidimensionality of touch, and underlining the need for comprehensive, modality-specific assessment of tactile imagery ability.

## INTRODUCTION

“Mental imagery is like perceiving, but in the absence of an immediate appropriate sensory stimulation” (Kosslyn et al. (2010, p. 3)). Mental imagery depends on stored and perceptually-grounded representations, involving specific sensory modalities and shows neural activation of sensory processing areas similar to that during perception of modality-specific external stimuli (Albers et al., 2013; McNorgan, 2012). Indeed, imagery and perception share important functional characteristics and can influence each other (Anema et al., 2012; Ishai & Sagi, 1995), even though their experience can be very different (Koenig-Robert & Pearson, 2020). There are also substantial individual differences in imagery ability (Kosslyn et al., 1984; Sulfaro et al., 2024). These are usually expressed in how vividly someone experiences their imagery (Lacey & Lawson, 2013), with on the extreme end people with aphantasia, who are not able to induce imagery (Zeman et al., 2015). Interestingly, aphantasia can be limited to one modality, suggesting that the ability to conjure a vivid image can differ between modalities (Arnold et al., 2026; Dawes et al., 2024). It can even differ across different dimensions *within* a modality (Palermo et al., 2022).

Most investigations of the functional and neural characteristics of imagery have concerned visual, motor and to some extent auditory modalities, while the tactile modality has received limited attention (Gallace, 2013; McNorgan, 2012). At the same time, the crucial role of touch for many aspects of our daily functioning, from exploring our environment, to guiding our action and providing social and emotional support, also has become increasingly clear (de Haan & Dijkerman, 2020). The limited interest in tactile imagery may also be related to the complexity and multidimensionality of the sense of touch. Several aspects of touch can be identified. An important distinction is that between passively receiving touch or haptically exploring objects and other items in the environment (Gibson, 1962; Prescott et al., 2011). While perceiving passive touch depends entirely on the different receptors in the skin, active touch involves spatiotemporal integration of input from the skin receptors with those in muscles, tendons and joints, while actively exploring an object, thus adding a motor component as well (Lepora, 2016). Active touch usually involves our hands and fingers and is focused on identifying different features such as texture, temperature, surface compliance and the weight of an object, with each feature requiring a specific exploratory procedure (Lederman & Klatzky, 1993). In contrast, passive touch can be received across different body parts, that vary widely in their tactile sensitivity (Corniani & Saal, 2020; Weinstein, 1968). Moreover, different aspects of the skin contact can be experienced, from pressure to vibration, stroking to pain.

Considering that imagery vividness can vary across different dimensions within a modality (O’ Dowd et al., 2022; Palermo et al., 2022), these multidimensional aspects of touch should be reflected in tactile imagery research as well. Studies of tactile imagery have focused on similarities to tactile stimulation either in terms of neural processing or regarding functional characteristics. With respect to neural processing related to active tactile imagery, Newman et al. (2005) reported a distinction between imagery of haptic material (roughness, hardness, temperature) and geometric object (shape, size) features. While imagining geometric features evoked visual imagery and resulted in activation in and around the intraparietal sulcus, imagining material features evoked processing of semantic object representations and resulted in activation in the inferior extrastriate areas. The neural basis of passive tactile imagery has been investigated more extensively, in particular whether the primary somatosensory cortex is involved (which it is, (Morozova et al., 2025; Schmidt et al., 2014)). Indeed, neural activity during imagery of touch at different body sites shows a somatotopic organisation similar to the neural activity in response to physical touch across different body sites (Kaas et al., 2019; Schmidt & Blankenburg, 2019). Moreover, multivariate pattern analysis shows that the neural activity in primary somatosensory cortex during imagined touch reflects the contents of that touch stimulus (Nierhaus et al., 2023). Further evidence for similarities between actual and imagined touch come from the hedonic experience of touch. Several studies have shown that the pattern of pleasantness ratings of different types of imagined tactile stroking is similar to those of physical tactile stroking (Lustenhouwer & Meijer, 2026; Panagiotopoulou et al., 2018; Tagini et al., 2023).

Taken together, these studies show that the multidimensionality of touch is reflected in tactile imagery, though most of them have focused on one aspect of active or passive touch imagery in isolation. Recently however, O’Dowd and colleagues (2022) explored different aspects of tactile imagery in a single study. They investigated the perceived vividness of imagining active touch (exploring objects haptically and judging the vividness of their texture, compliance, force and weight) and sensitivity of imagined passive tactile input (being stroked with a brush on various body sites). They reported object- and property-specific differences in vividness for imagining active touch and varying sensitivity for imagined stroking across body sites. Interestingly, the sensitivity of imagined stroking followed a pattern similar to tactile sensitivity across these body sites: imagery was generally more sensitive for body sites with higher tactile sensitivity (i.e. with lower two-point discrimination thresholds and higher receptor density). Finally, the authors reported substantial individual variability and vividness of active tactile imagery and sensitivity of passive tactile imagery were only weakly correlated, further highlighting individual differences and the multidimensional nature of tactile imagery.

Although current literature suggests that tactile imagery shares functional and neural characteristics with tactile perception, is similarly multidimensional and can vary depending on the object explored or the touched body site, several important aspects remain to be explored. First, most studies, like O’Dowd et al. (2022), have focused solely on vividness of tactile imagery. Vividness is thought to capture how well an individual can *generate* and *inspect* an imagined sensation (Kosslyn, 1995; Kosslyn et al., 1984). As such, it captures only two of the four core imagery processing components as defined by Kosslyn and colleagues (1984). It is unclear how well individuals can *maintain* and *transform* imagined touch. Second, imagery of different types of passive tactile sensations (e.g. stroking, vibration, pinching) has only been studied separately. Since a direct comparison within one study is lacking, it is unclear whether imagery characteristics such as vividness, maintenance and transformation vary depending on the type of somatosensory sensation. The current study therefore aims to map a comprehensive assessment of tactile imagery in terms of imagery processing components (vividness, maintenance, transformation), type of touch (active versus passive) and touch qualities (object properties for active touch, different tactile sensations across body sites for passive touch). In addition, we assess experienced pleasantness in a subset of participants, as hedonic experience, an important aspect of touch perception (McGlone et al., 2014), is similar for tactile imagery and stimulation, and a link between stimulus affect and imagery vividness has been reported in other imagery modalities (Bywaters et al., 2004; Cui et al., 2025). Ultimately, this investigation might lead to the development of a comprehensive tactile imagery ability assessment tool, something which current literature lacks.

While this study is exploratory in nature, there are some general expectations: First, based on the O’Dowd (2022) study, we anticipate imagery ability (quantified as its vividness, maintenance and transformation) to vary between object properties and body sites. With respect to the latter, we anticipate higher ability for more sensitive body sites (i.e. the lip and sole of the foot) compared to less sensitive body sites (i.e. the lower back and the shin). Second, also based on O’Dowd, we anticipate that imagery ability for passive and active touch are related to some extent, but do not fully overlap. We expect a similar relation between the different imagery processing components, as Kosslyn’s model (1984) posits that they capture overlapping, yet distinct processes. Fourth, since hedonic experience of tactile imagery mirrors that of tactile stimulation, imagery of painful sensations, in this case being pinched, is expected to be experienced as less pleasant than neutral or positive sensations like vibration and stroking. Finally, given the relation between affect and imagery vividness in other imagery modalities (Bywaters et al., 2004; Cui et al., 2025) we expect that imagery ability relates to experienced pleasantness.

## METHODS

This study was reviewed and approved by the Ethics Review Board of Utrecht University’s Faculty of Social and Behavioural Sciences.

### Participants

All participants were at least 18 years of age and had normal or corrected-to-normal vision. Synesthesia, or presence of any cognitive, neurological, somatosensory and/or tactile impairments were reason for exclusion. Participants were recruited through the Faculty of Social and Behavioural Sciences’ research participation system (SONA) and convenience sampling. Students from the undergraduate Psychology program could receive research participation credits for their participation.

### Procedure

Data collection occurred between December 2024 and February 2026 in the Faculty’s research laboratories or in a quiet setting off-campus. Data were recorded on a laptop using Qualtrics survey software (Provo, UT). After giving informed consent, participants sat down at a table and provided their demographic information (age, gender). Trained assessors then instructed participants to imagine different active and passive tactile sensations and recorded participant’s imagery vividness, maintenance and transformation. A subset of participants also rated imagery pleasantness. Finally, participants answered five additional questions about their experience with the task and tactile imagery in general (outside the scope of this article, see suppl. materials).

### Design

#### Tactile imagery ability

For this study, we developed a tool to assess tactile imagery ability, quantified as imagery vividness, maintenance and transformation (see suppl materials), which consisted of two sections: active and passive tactile imagery, totaling 72 items. Participants were instructed to keep their eyes closed during the imagery and to focus on the feeling of touch, while refraining from using visual information.

The active (haptic) section consisted of 36 items, in which participants were asked to imagine the sensation of touching different objects and their properties. We included three objects (plastic bottle, modelling clay, sponge) and four properties (surface resistance, weight, texture, temperature). We selected objects for which transformation of all four properties was feasible. For each object-property combination, participants were first instructed to imagine that specific sensation, after which they rated its vividness on a 5-point Likert scale, ranging from 1 (*No imagery - I only know that I am thinking about the sensation*) to 5 (*Very detailed and vivid – the imagery of the sensation is as vivid as a real stimulus*). For bottle-weight for example, participants were instructed to *‘Imagine the strength needed to lift an empty water bottle.’* and rate ‘*How vividly can you imagine the sensation of its weight?’*. They were then asked to imagine the same sensation, hold it for 5 seconds, and rate how well they were able to maintain the sensation on a 5-point Likert scale, ranging from 1 (*Unable to maintain the imagery of the sensation – it fades quickly and cannot be sustained*) to 5 (*Completely able to maintain the imagery of the sensation – it is steady and does not fade over time*). Finally, they were instructed to transform the sensation and rate how well they were able to control the instructed change on a 5-point Likert scale ranging from 1 (*Unable to control the sensation – I cannot manipulate or change the imagery of the sensation at all*) to 5 (*Completely able to control the sensation – I can fully manipulate or change the imagery of the sensation as intended.*). In the bottle-weight example, they were instructed to ‘*Now imagine the bottle is gradually getting heavier as it fills up with water.*’ and rate ‘*How well are you able to control the change of this sensation?’.* Instructed transformations went from softer to harder (clay, sponge) or vice versa (bottle) for surface resistance, from lighter to heavier for weight, from smoother to rougher for texture and from colder to warmer for temperature.

The passive (somatosensory) section also consisted of 36 items. Here participants were asked to imagine the sensation of being touched at different body sites. We included four body sites (lower lip, sole of the right foot, left shin, lower back), and three different tactile sensations (stroking, pinching, vibration). Body sites were selected to include both non-glabrous and glabrous skin sites, with different levels of receptor density (Corniani & Saal, 2020; Weinstein, 1968) and imagery sensitivity (O’ Dowd et al., 2022). Participants were first instructed to imagine a specific sensation on a specific body site and rate its vividness on the same 5-point Likert scale as in the active part (ranging from 1 *– No imagery,* to 5 – *Very detailed and vivid*). For example, for the lower back-vibration combination, the instructions were to *‘Imagine the feeling of a massage gun lightly vibrating on your lower back’* and to rate ‘*How vividly can you imagine this sensation?*’. They were then instructed to transform that sensation and rate the level of control over the instructed change on the 5-point Likert scale for transformation (ranging from 1 – *Unable to control the sensation* to 5 – *Completely able to control the sensation*). In the back-vibration example, they were instructed to ‘*Imagine that the light vibration of the massage gun on your lower back gradually increases in pressure, making the vibration harder*.’ and rate ‘*How well are you able to control the change of this sensation?* Transformation instructions were a gradual increase in stroking pressure starting from light/gentle stroking, a gradual increase in pinching pressure going from slightly painful to more painful, and a gradual increase in vibration pressure (for the foot, shin and back). For the lip, vibration changed from light and constant to pulsating. Finally, participants were instructed to imagine the untransformed sensation again, hold it for 5 seconds, and rate how well they were able to maintain the sensation on the 5-point Likert scale for maintenance (ranging from 1 - *Unable to maintain the imagery of the sensation* to 5 - *Completely able to maintain the imagery of the sensation*).

#### Pleasantness of tactile imagery

A subset of 46 participants additionally rated the pleasantness of all object-property and body site-sensation pairs. After each pair’s three imagery ability items (vividness, maintenance, transformation), they indicated how pleasant or unpleasant they found the imagery experience, by placing a slider along a Visual Analogue Scale (VAS) ranging from ‘*unpleasant*’ (-10) to ‘*pleasant*’ (10). The scale’s midpoint (0) was labeled ‘*neutral*’. Pleasantness ratings were rounded to the nearest integer by Qualtrics at the time of recording.

### Analyses

Since imagery vividness, maintenance and transformation capture different imagery processing components (Kosslyn et al., 1984) and were measured using different scales, we conducted separate analyses for each of them.

Data were first exported from Qualtrics as .CSV files, which were then imported in Python (version 3.13). We calculated mean imagery ability and pleasantness scores for active and passive tactile imagery by collapsing over all conditions. We also calculated mean imagery ability and pleasantness scores for each condition of interest.

### Statistical analyses

All pairwise comparisons were conducted in JASP version 0.19.1 (JASP Team, 2024). All correlations were conducted in Python, using SciPy’s *stats* package (Virtanen et al., 2020). Where appropriate, we applied Holm-Bonferroni corrections for multiple comparisons (Holm, 1979), using the *multipletests* function from the statsmodels package in Python (Seabold & Perktold, 2010).

All factorial ANOVAs (vividness, maintenance, transformation, pleasantness) and subsequent post-hoc testing were also conducted in JASP. If Mauchly’s testing indicated violation of the assumption of sphericity, we applied appropriate corrections (Greenhouse-Geisser (if ε < .75) or Huynh-Feldt (H-F) (if ε > .75) (Girden, 1992). If main or interaction effects warranted post-hoc testing, we applied JASP’s build-in Holm-Bonferroni corrections (Holm, 1979).

#### Tactile imagery ability

We explored differences between active and passive tactile imagery through pairwise comparisons of active and passive imagery ability scores. Since the assumption of normality was violated for vividness and transformation, we conducted Wilcoxon signed-rank tests. We compared active and passive maintenance scores with a paired-samples t-test. We applied a Holm-Bonferroni correction (n=3).

To test the relation between active and passive tactile imagery ability, we conducted three Spearman correlations using the *spearmanr* function on each of the imagery processing components. Here we also corrected for multiple comparisons (n=3) through a Holm-Bonferroni correction.

We also explored the relation between the different imagery processing components by correlating overall vividness, maintenance and transformation with each other, separately for active and passive tactile imagery. We used the *pearsonr* function to conduct a Pearson correlation for active imagery maintenance vs. transformation, and *spearmanr* for all other correlations. Holm-Bonferroni corrections were applied for active and passive imagery (n=3 each).

To explore how active tactile imagery ability varies across objects and their properties, we conducted three separate 3×4 factorial ANOVAs with factors OBJECT (bottle, clay, sponge) and PROPERTY (resistance, weight, texture, temperature) on imagery vividness, maintenance, and transformation scores.

We investigated how passive tactile imagery ability varies across body sites and tactile sensations, by running three 4×3 factorial ANOVAs with factors BODY SITE (lip, foot, shin, back) and SENSATION (stroking, pinching, vibration), again on imagery vividness, maintenance, and transformation scores.

#### Imagery pleasantness

We compared overall pleasantness of active and passive tactile imagery with a Wilcoxon signed-rank test, as data were not normally distributed. We also explored the relation between pleasantness of active and passive tactile imagery by calculating Spearman’s correlations on active and passive pleasantness scores.

To explore experienced pleasantness during active and passive tactile imagery, we conducted two separate factorial ANOVAs on pleasantness scores: a 3×4 ANOVA (OBJECT x PROPERTY) for active, and a 4×3 ANOVA (BODY SITE x SENSATION) for passive tactile imagery.

#### Correlations between imagery ability and pleasantness

To explore the relation between hedonic experience and imagery ability, we correlated overall mean pleasantness with overall mean ability for active tactile imagery. Since pleasantness scores differed per sensation for passive tactile imagery, we correlated pleasantness and imagery ability scores per sensation (stroking, pinching, vibration). When data were normally distributed, we calculated Pearson correlations with the *pearsonr* function. When the assumption of normality was violated, we used the *spearmanr* function to correlate mean pleasantness scores with mean imagery vividness, maintenance and transformation. We applied Holm-Bonferroni corrections (n=3) for both active and passive correlations.

## RESULTS

### Participants characteristics

The full sample (n =136) included 91 women, 44 men and 1 other gender (mean age = 24±5.5 years). Pleasantness scores were obtained in a subsample (n = 46) of 34 women and 12 men (mean age = 23±6 years).

### Tactile imagery ability

Mean scores for both active and passive tactile imagery indicate that participants were generally able to perform tactile imagery relatively well (see Table 1). Overall, imagery was rated as at least *relatively clear and vivid* (3 or higher) by 84% for active and 80% for passive tactile imagery. Likewise, the ability to maintain and transform the imagery were rated as at least *moderate* (3 or higher) by 63% and 77% for active and 62% and 71% for passive imagery respectively. Individual variability was substantial, as score ranges spanned both ends of the scale (Table 1, Fig.1,). There were no significant differences in ability for active compared to passive tactile imagery (Table 1).

**Figure 1.**
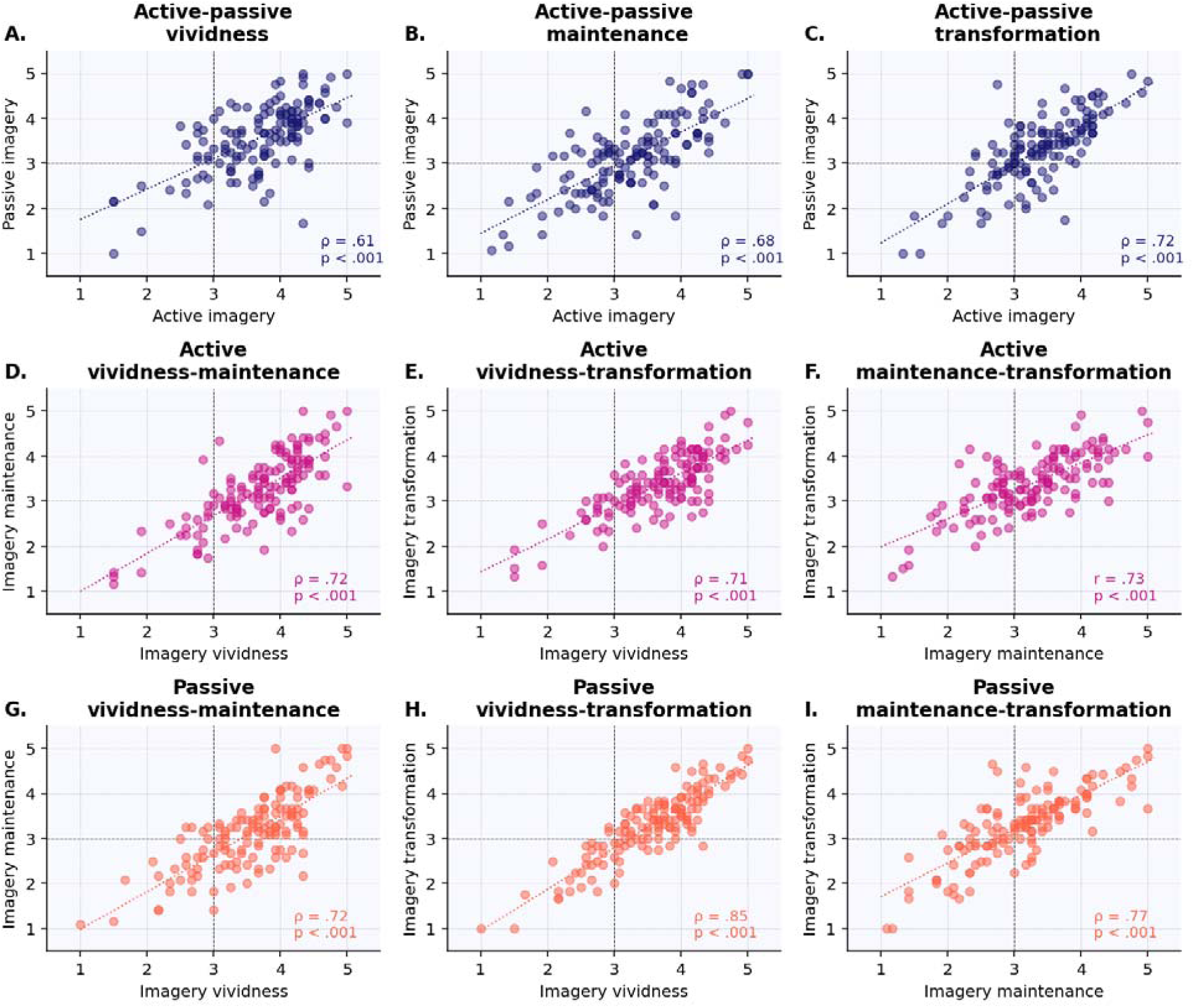
Correlations between imagery ability scores measured on 5 point Likert scales. Higher scores indicate higher ability. A.-C. show correlations between active and passive imagery, for vividness, maintenance and transformation. D.-F. show correlations within active tactile imagery, between imagery processing components, vividness and maintenance (D.), vividness and transformation (E.) and maintenance and transformation (F.). G.-I. show correlations within passive tactile imagery, between imagery processing components, vividness and maintenance (G.), vividness and transformation (H.) and maintenance and transformation (I.). Dotted black lines denote scale midpoints (3). Dotted colored lines are trendlines. p = p_holm_, Holm-Bonferroni corrected p-value (n=3); ρ = Spearman’s rho; r = Pearson’s r.

**Table 1.** Overall active and passive imagery vividness, maintenance, transformation and pleasantness. Imagery ability is measured on 5-point Likert scales, with higher scores indicating higher imagery ability. Pleasantness is measures on a Visual Analogue Scale ranging from -10 (unpleasant) to 10 (pleasant). Descriptives, and statistics for pairwise comparisons and correlations between active and passive tactile imagery. d = Cohen’s d; IQR = interquartile range; max = maximum score; min = minimum score; SD = standard deviation; pholm = Holm-Bonferroni corrected p-value adjusted for comparing a family of 3; r = Pearson’s r; rB = rank-biserial correlation coefficient; ρ = Spearman’s rho; VAS = Visual Analogue Scale; W = Wilcoxon’s test statistic.

|  | Active<br>imagery | Passive<br>imagery | Pairwise<br>comparison<br>active-passive | Correlation<br>active-passive |
| --- | --- | --- | --- | --- |
| **Imagery ability (n=136)** |  |  |  |  |
| **Vividness** |  |  |  |  |
| mean ± SD | 3.68 ± 0.70 | 3.56 ± 0.74 | W(135) = 4797, | ρ(134) = .61, |
| median, IQR | 3.75, 0.92 | 3.67, 1.00 | p<sub>holm</sub> = .068, r<sub>B</sub> =<br>.22 | p<sub>holm</sub> < .001 |
| min-max | 1.5-5 | 1-5 |  |  |
| **Maintenance** |  |  |  |  |
| mean ± SD | 3.25 ± 0.78 | 3.13 ± 0.81 | t(135) = 2.32, | r(134) = .68, |
| median, IQR | 3.25, 1.02 | 3.17, 1.00 | p<sub>holm</sub> = .066, d =<br>0.20 | p<sub>holm</sub> < .001 |
| min-max | 1.17-5 | 1.08-5 |  |  |
| **Transformation** |  |  |  |  |
| mean ± SD | 3.38 ± 0.66 | 3.31 ± 0.77 | W(135) = 730, | ρ(134) = .72, |
| median, IQR | 3.42, 0.85 | 3.42, 0.94 | p<sub>holm</sub> = .068, r<sub>B</sub> =<br>.20 | p<sub>holm</sub> < .001 |
| min-max | 1.33-5 | 1-5 |  |  |
| **Pleasantness (VAS)**<br>**(n=46)** |  |  |  |  |
| mean ± SD | 1.61 ± 2.25 | 0.37 ± 2.13 | W(45) = 730 | ρ(44) = .11, |
| median, IQR | 1.33, 2.25 | 0.54, 2.42 | $p = .006$, $r_B = .48$ | $p = .480$ |
| min-max | -7.58-6.67 | -6.92-5.92 |  |  |

#### Correlation between active and passive tactile imagery ability

Three separate Spearman correlations revealed significant positive correlations between active and passive tactile imagery for each of the imagery processing components. Individuals with more vivid *active* tactile imagery also had more vivid *passive* tactile imagery (Table 1, Fig.1A.). Likewise, individuals with higher maintenance and transformation during *active* tactile imagery also showed higher maintenance and transformation during *passive* imagery (Table 1, Fig.1B.-C.).

#### Correlation between tactile imagery processing components

To explore the relation between the different imagery processing components, we correlated imagery vividness, maintenance and transformation with each other, separately for active and passive imagery. For both active and passive tactile imagery, all three imagery processing components were significantly positively correlated with each other (p_holm_ < . 001), with large effect sizes (see Fig1D.-I. for exact effect sizes).

#### Active tactile imagery

To understand how active tactile imagery varies across different types of active touch, we investigated imagery vividness, maintenance, and transformation across several objects and their properties.

##### Vividness

Vividness of active tactile imagery varied significantly across objects (OBJECT: F(2,239) = 6.25, p = .003, η_p_^2^ = .04, H-F corrected, Fig.2A.), but not significantly across properties (PROPERTY: F(3,361) = 2.62, p = .057, η_p_^2^ = .02, H-F corrected, Fig.2B.). Post-hoc testing revealed that overall vividness was significantly higher for the bottle than the clay (t(135)= 3.19, p_holm_ = .005, d = 0.20) and the sponge (t(135)= 2.43, p_holm_ = .033, d = 0.10). Vividness of the sponge and the clay did not differ significantly (t(135)= 1.50, p_holm_ = .14, d = 0.08).

**Figure 2.**
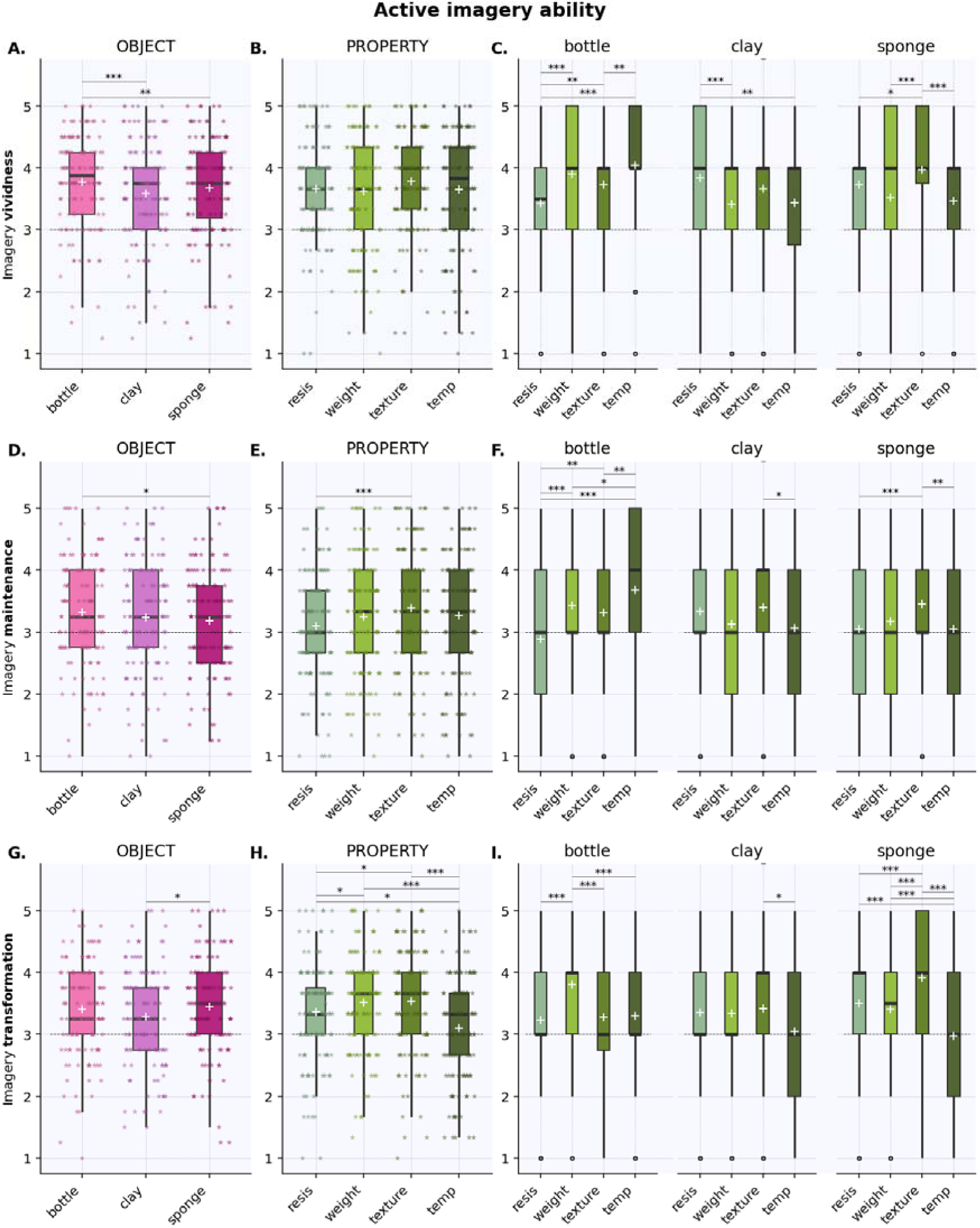
ACTIVE tactile imagery ability of objects and their properties. Plots show main effects of OBJECT (A., D., G.) and PROPERTY (B., E., H.) and the interaction between them (C.,F.,I.) for active imagery vividness (A.-C.), maintenance (D.-F.) and transformation (G.-I.). Scores range from 1 to 5, with higher scores denoting higher imagery ability. Boxes represent the interquartile ranges (IQR (from the 25^th^ (Q1) to the 75^th^ (Q3) percentile); whiskers show the minimum within Q1 -1.5 times the IQR and maximum within Q3 + 1.5 times the IQR. The black lines show the median, and the white + the mean per condition. The dashed line in each plot denotes the scale midpoint (3). = individual data points, jittered for visibility purposes; ◦ **=** outlier; * significant difference at p_holm_ ≤.05; ** significant difference at p_holm_ ≤.01; *** significant difference at p_holm_ ≤.001. resis = resistance; temp = temperature.

There was a significant interaction between OBJECT and PROPERTY (F(6,763) = 15.66, p < .001, η_p_^2^ = .10, H-F corrected), indicating that imagery vividness of specific properties varied per object. We followed-up this effect through conditional comparisons based on factor OBJECT (Fig. 2C, statistics in Table 2). For the bottle, resistance was significantly *less* vivid than all other properties, and texture was significantly *less* vivid than temperature. The clay showed a reversed pattern where resistance was significantly *more* vivid than temperature and weight. The sponge showed a similar trend of *more* vivid resistance than temperature. In further contrast with the bottle, the sponge’s texture was significantly *more* vivid than its temperature and weight. The clay showed a similar trend, for texture being more vivid than weight. Similar to the bottle, the sponge showed significantly *less* vivid resistance than texture.

**Table 2.**
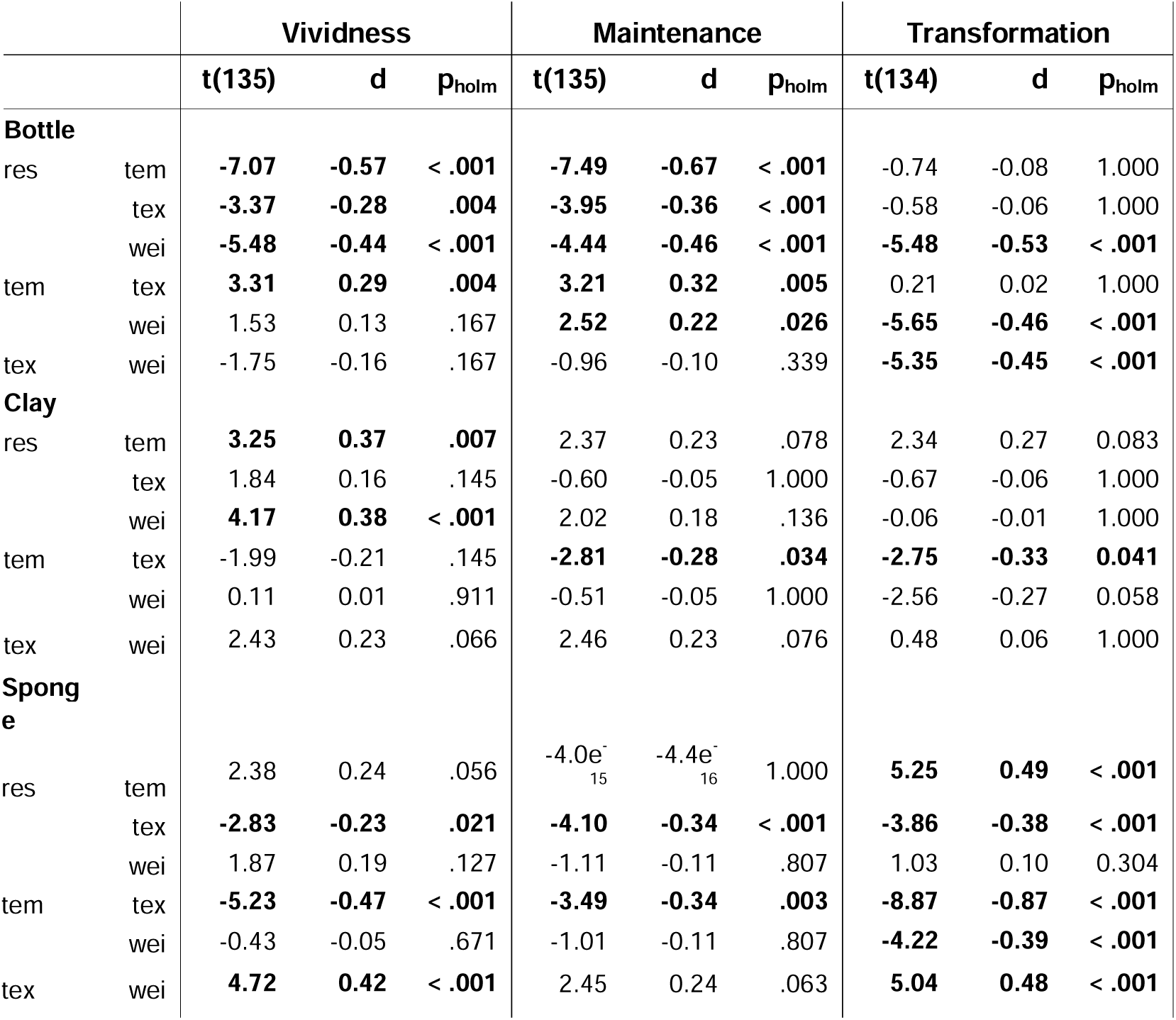
Post-hoc pairwise comparisons for OBJECT x PROPERTY interaction effects for active tactile imagery ability (vividness, maintenance, and transformation). Conditional on factor OBJECT. d = Cohen’s d; pholm = Holm-Bonferroni corrected p-value adjusted for comparing a family of 6; res = surface resistance; tem = temperature; tex = texture; wei = weight. Significant differences are printed in bold.

|  |  | Vividness |  |  | Maintenance |  |  | Transformation |  |  |
| --- | --- | --- | --- | --- | --- | --- | --- | --- | --- | --- |
|  |  | t(135) | d | p<sub>holm</sub> | t(135) | d | p<sub>holm</sub> | t(134) | d | p<sub>holm</sub> |
| **Bottle** |  |  |  |  |  |  |  |  |  |  |
| res | tem | **-7.07** | **-0.57** | **< .001** | **-7.49** | **-0.67** | **< .001** | -0.74 | -0.08 | 1.000 |
|  | tex | **-3.37** | **-0.28** | **.004** | **-3.95** | **-0.36** | **< .001** | -0.58 | -0.06 | 1.000 |
|  | wei | **-5.48** | **-0.44** | **< .001** | **-4.44** | **-0.46** | **< .001** | **-5.48** | **-0.53** | **< .001** |
| tem | tex | **3.31** | **0.29** | **.004** | **3.21** | **0.32** | **.005** | 0.21 | 0.02 | 1.000 |
|  | wei | 1.53 | 0.13 | .167 | **2.52** | **0.22** | **.026** | **-5.65** | **-0.46** | **< .001** |
| tex | wei | -1.75 | -0.16 | .167 | -0.96 | -0.10 | .339 | **-5.35** | **-0.45** | **< .001** |
| **Clay** |  |  |  |  |  |  |  |  |  |  |
| res | tem | **3.25** | **0.37** | **.007** | 2.37 | 0.23 | .078 | 2.34 | 0.27 | 0.083 |
|  | tex | 1.84 | 0.16 | .145 | -0.60 | -0.05 | 1.000 | -0.67 | -0.06 | 1.000 |
|  | wei | **4.17** | **0.38** | **< .001** | 2.02 | 0.18 | .136 | -0.06 | -0.01 | 1.000 |
| tem | tex | -1.99 | -0.21 | .145 | **-2.81** | **-0.28** | **.034** | **-2.75** | **-0.33** | **0.041** |
|  | wei | 0.11 | 0.01 | .911 | -0.51 | -0.05 | 1.000 | -2.56 | -0.27 | 0.058 |
| tex | wei | 2.43 | 0.23 | .066 | 2.46 | 0.23 | .076 | 0.48 | 0.06 | 1.000 |
| **Sponge** |  |  |  |  |  |  |  |  |  |  |
| res | tem | 2.38 | 0.24 | .056 | -4.0e<sup>-15</sup> | -4.4e<sup>-16</sup> | 1.000 | **5.25** | **0.49** | **< .001** |
|  | tex | **-2.83** | **-0.23** | **.021** | **-4.10** | **-0.34** | **< .001** | **-3.86** | **-0.38** | **< .001** |
|  | wei | 1.87 | 0.19 | .127 | -1.11 | -0.11 | .807 | 1.03 | 0.10 | 0.304 |
| tem | tex | **-5.23** | **-0.47** | **< .001** | **-3.49** | **-0.34** | **.003** | **-8.87** | **-0.87** | **< .001** |
|  | wei | -0.43 | -0.05 | .671 | -1.01 | -0.11 | .807 | **-4.22** | **-0.39** | **< .001** |
| tex | wei | **4.72** | **0.42** | **< .001** | 2.45 | 0.24 | .063 | **5.04** | **0.48** | **< .001** |

To further explore the interaction effect, we additionally performed conditional comparisons based on factor PROPERTY (Supl. Table 1, Supl. Fig. 1). This revealed that imagery of the bottle was significantly *less* vivid than the clay and the sponge for resistance, whereas it was significantly *more* vivid than the clay and the sponge for both temperature and weight. Finally, the sponge was significantly *more* vivid than the bottle and the clay for texture.

##### Maintenance

As with vividness, there was a significant main effect of OBJECT for maintenance of active tactile imagery (F(2,270) = 3.24, p = .041, η_p_^2^ = .02, Fig.2D). Post-hoc testing revealed that maintenance of the bottle was significantly higher than that of the sponge (t(135)= 2.73, p_holm_ = .022, d = 0.12). Imagery maintenance did not differ significantly between the bottle and the clay (t(135)= 1.54, p_holm_ = .251, d = 0.08) nor the clay and the sponge (t(135)= 0.84, p_holm_ = .403, d = 0.04).

The main effect of PROPERTY was also significant for maintenance (F(3,405) = 5.63, p < .001, η_p_^2^ = .04, Fig.2E.), indicating that ability to maintain imagery depended on the specific property. Post-hoc testing revealed that maintenance of resistance was significantly lower than that of texture (t(135)= -4.62, p_holm_ < .001, d = -0.25). There was also a trend for lower maintenance of resistance compared to temperature (t(135)= -2.44, p_holm_ = .080, d = -0.15). The other properties did not differ significantly from each other (Suppl. Table 2).

Similar to vividness, OBJECT and PROPERTY interacted significantly (F(6,810) = 11.57, p < .001, η_p_^2^ = .08), indicating that imagery maintenance varied depending on specific object-property combinations. We conducted post-hoc conditional comparisons based on factor OBJECT (Fig. 2F, statistics in Table 2). For the bottle, maintenance of temperature was significantly *higher* than all other properties. Maintenance of bottle resistance was also significantly *lower* than that of texture and weight. The clay showed a reversed pattern, where maintenance was significantly *lower* for temperature than texture. There were also trends for lower maintenance of clay’s temperature than resistance, and for higher maintenance of clay’s texture than weight. Finally, maintenance of the sponge was significantly *lower* for resistance than for texture, similar to the bottle. In contrast with the bottle, but similar to the clay, maintenance of the sponge was significantly *higher* for texture compared to temperature. As with the clay, there was a trend for higher maintenance of the sponge’s texture than its weight.

Additional conditional comparisons based on factor PROPERTY (Suppl.Table3, Suppl.Fig.2) showed that, similar to vividness, the bottle had significantly *higher* maintenance than clay and sponge for temperature and weight, but significantly *lower* maintenance than the clay for resistance. Unlike vividness, maintenance of resistance was significantly higher for the clay than for the sponge. Imagery maintenance of texture did not differ significantly between objects.

##### Transformation

In line with vividness and maintenance, there was a significant main effect of OBJECT for transformation of active tactile imagery (F(2,256) = 4.09, p = .018, η_p_^2^ = .03, H-F corrected, Fig.2G.). Post-hoc tests revealed that overall imagery transformation was significantly lower for the clay than for the sponge (t(134)= -2.66, p_holm_ = .026, d = -0.14). Transformation did not differ significantly between remaining objects (Suppl. Table 4).

As with maintenance, imagery transformation varied significantly across properties (F(3,402) = 18.59, p < .001, η_p_^2^ = .12, Fig.2H.). Post-hoc tests showed that overall transformation of temperature was significantly lower than transformation of all other properties (resistance: t(134)= -3.64, p_holm_ = .002, d = -0.23; texture: t(134)= -6.25, p_holm_ < .001, d = -0.39; weight: t(134)= -7.01, p_holm_ < .001, d = -0.37). Transformation of resistance was significantly lower than texture (t(134)= -2.93, p_holm_ = .012, d = -0.17) and weight (t(134)= -2. 64, p_holm_ = .030, d = -0.15). Transformation of texture and weight did not differ significantly (t(134)= 0.30, p_holm_ = .766, d = 0.02).

Similar to vividness and maintenance, imagery transformation showed a significant OBJECT x PROPERTY interaction effect (F(6,776) = 11.52, p < .001, η_p_^2^ = .08, H-F corrected). Post-hoc conditional comparisons based on factor OBJECT revealed that imagery transformation differed between object-property pairs (Fig.2I, statistics in Table 2). For the bottle, transformation of *weight* was significantly higher than all other properties. In contrast, for the sponge, transformation of *texture* was significantly higher than all other properties. Moreover, transformation of sponge temperature was significantly lower than that of sponge resistance and weight. The clay showed a partially similar pattern, where transformation of temperature was significantly lower than texture. The clay also showed trends towards lower transformation of temperature than resistance and weight.

Additional conditional comparisons based on factor PROPERTY (Suppl.Table.5, Suppl.Fig.3) showed that transformation of resistance and texture was significantly *higher* for the sponge than for the bottle. The opposite pattern emerged for temperature and weight, where transformation was significantly *lower* for the sponge compared to the bottle. Transformation of the clay was significantly lower than the sponge for texture, and significantly lower than the bottle for weight. For temperature, transformation of the clay showed a trend towards being lower than the bottle, similar to the clay.

**Figure 3.**
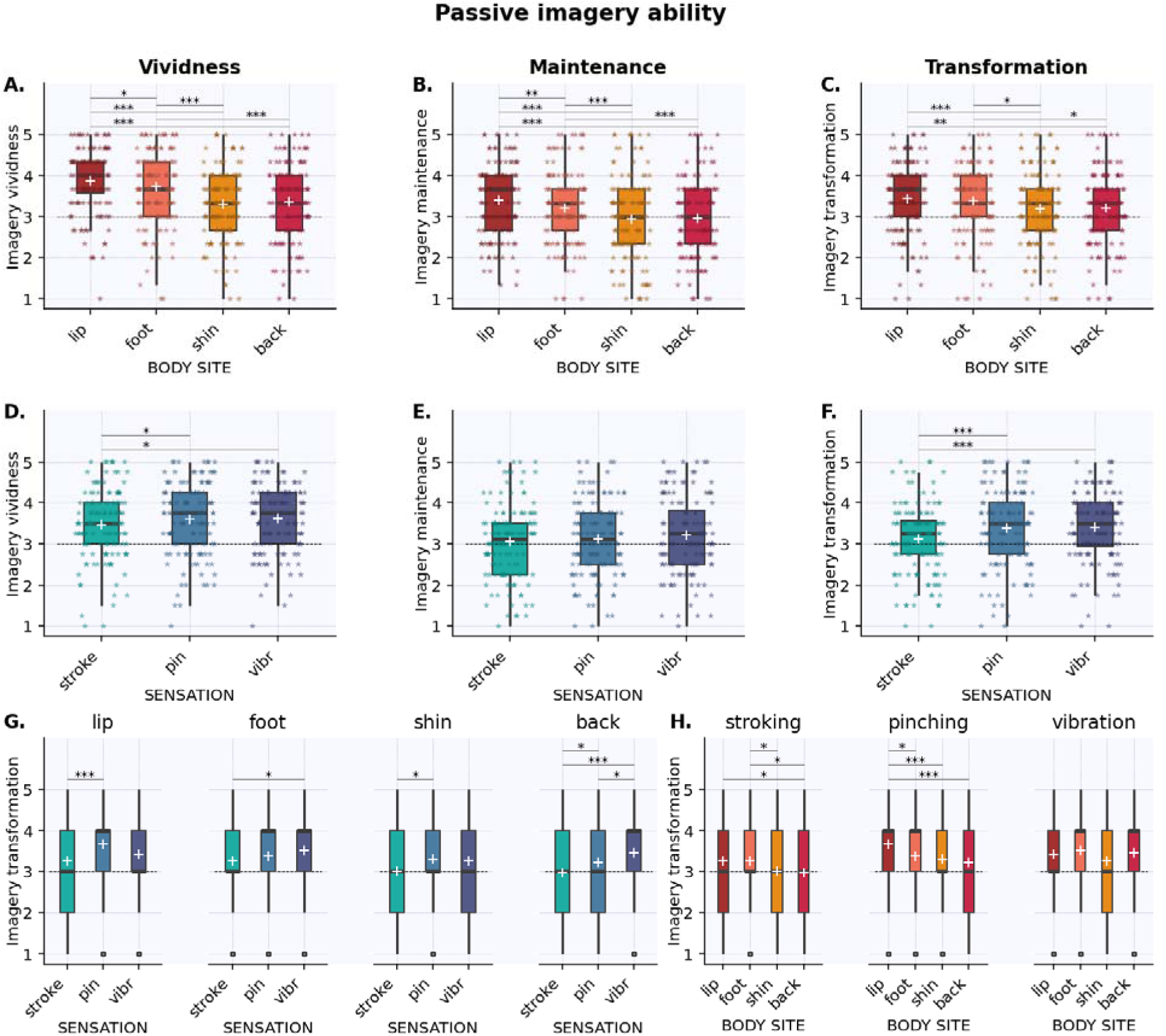
PASSIVE tactile imagery ability of different sensations across body sites. Plots show main effects of BODY SITE (A.-C) and SENSATION (D.-F.) for vividness (A.,D.), maintenance (B.,E.), and transformation (C.,F.), and the significant interaction between them for transformation (G.,H.). Scores range from 1 to 5, with higher scores denoting higher imagery ability. Boxes represent the interquartile ranges (IQR (from the 25^th^ (Q1) to the 75^th^ (Q3) percentile); whiskers show the minimum within Q1 -1.5 times the IQR and maximum within Q3 + 1.5 times the IQR. The black lines show the median, and the white + the mean per condition. The dashed line in each plot denotes the scale midpoint (3). = individual data points, jittered for visibility purposes; ◦ **=** outlier; * significant difference at pholm ≤.05; ** significant difference at pholm ≤.01; *** significant difference at pholm ≤.001. stroke = stroking; pin = pinching; vibr = vibration.

#### Passive tactile imagery

To investigate how tactile imagery varies across different types of passive touch, we tested how imagery vividness, maintenance, and transformation varied across body sites and various tactile sensations.

##### Vividness

Vividness of passive tactile imagery showed a main effect of BODY SITE (F(3,405) = 37.67, p < .001, η_p_^2^ = .22, Fig.3A.). Post-hoc tests revealed that imagery on the lip was significantly more vivid than all other body sites (foot: t(135) = 2.55, p_holm_ = .024, d = 0.13; shin: t(135) = 9.46, p_holm_ < .001, d = 0.52; back: t(135) = 8.06, p_holm_ < .001, d = 0.46). The foot was more vivid than the shin (t(135) = 6.22, p_holm_ < .001, d = 0.38) and the back (t(135) = 5.31, p_holm_ < .001, d = 0.33). Vividness did not differ between the shin and back (t(135) = -0.95, p_holm_ = .346, d = -0.06).

Imagery vividness also varied significantly across passive tactile sensations (F(2,270) = 3.54, p = .030, η_p_^2^ = .03, Fig.3D.). Post-hoc pairwise comparisons revealed that stroking was significantly less vivid than pinching (t(135) = 2.31, p_holm_ = .048, d = -0.12) and vibration (t(135) = -2.44, p_holm_ = .048, d = -0.13). Pinching and vibration did not differ in terms of vividness (t(135) = -0.18, p_holm_ = .861, d = -0.01).

BODY SITE and SENSATION did not interact significantly (F(6,810) = 1.06, p = .388, η_p_^2^ = .01) indicating that the different sensations showed similar patterns of vividness across body sites.

##### Maintenance

Imagery maintenance differed significantly across body sites (F(3,402) = 23.55, p < .001, η_p_^2^ = .15, Fig3B.). Similar to vividness, maintenance was significantly higher for the lip than all other body sites (foot: t(134) = 3.25, p_holm_ = .003, d = 0.17; shin: t(134) = 6.47.46, p_holm_ < .001, d = 0.40; back: t(134) = 6.64, p_holm_ < .001, d = 0.39). Maintenance for the foot was also significantly higher than for both the shin (t(134) = 4.26, p_holm_ < .001, d = 0.23) and the back (t(134) = 4.15, p_holm_ < .001, d = 0.22). Again, there was no significant difference between the shin and back (t(134) = -0.15, p_holm_ = .879, d = -0.01)

Imagery maintenance did not vary significantly across tactile sensations (F(2,268) = 2.84, p = .060, η_p_^2^ = .02, Fig.3E.), and there was no significant interaction between BODY SITE and SENSATION ((F(6,754) = 1.37, p = .227, η_p_^2^ = .01, H-F corrected).

##### Transformation

Imagery transformation differed significantly across body sites (F(3,405) = 7.73, p < .001, η_p_^2^ = .05, Fig.3C.). Post-hoc pairwise comparisons revealed a pattern mostly similar to vividness and maintenance, where transformation was significantly higher for the lip than for the shin (t(135) = 3.98, p_holm_ < .001, d = 0.23) and the back (t(135) = 3.50, p_holm_ = .003, d = 0.20). However, transformation of the lip and the foot did not differ significantly (t(135) = 1.12, p_holm_ = .527, d = 0.06). The foot again scored significantly higher than the shin (t(135) = 3.21, p_holm_ = .007, d = 0.17) and the back (t(135) = 2.45, p_holm_ = .046, d = 0.14). As with vividness and maintenance, transformation did not differ significantly between shin and back (t(135) = -0.46, p_holm_ = .648, d = -0.03).

As with vividness, imagery transformation varied across passive tactile sensations (F(3,405) = 12.56, p < .001, η_p_^2^ = .09, Fig.3F.) Transformation of stroking was significantly lower than both pinching (t(135) = -4.12, p_holm_ < .001, d = -0.23) and vibration (t(135) = -4.64, p_holm_ < .001, d = -0.25). Transformation of pinching and vibration did not differ significantly (t(135) = -0.31, p_holm_ = .759, d = -0.02).

There was a significant interaction between BODY SITE and SENSATION (F(6,810) = 2.51, p = .020, η_p_^2^ = .02), indicating that transformation of passive tactile imagery varied across body site-sensation pairs. We first explored the interaction through conditional comparisons on factor BODY SITE (Fig.3G, statistics in Table 3). Transformation of pinching was significantly higher than stroking for all body sites, except for the foot. Transformation of vibration was significantly higher than stroking for the foot and the back and showed a trend in the same direction for the shin. The back and the lip showed opposite patterns for transformation of pinching and vibration. Whereas transformation of vibration was significantly *higher* than pinching for the back, there was a trend towards *lower* transformation of vibration than pinching for the lip.

**Table 3.**
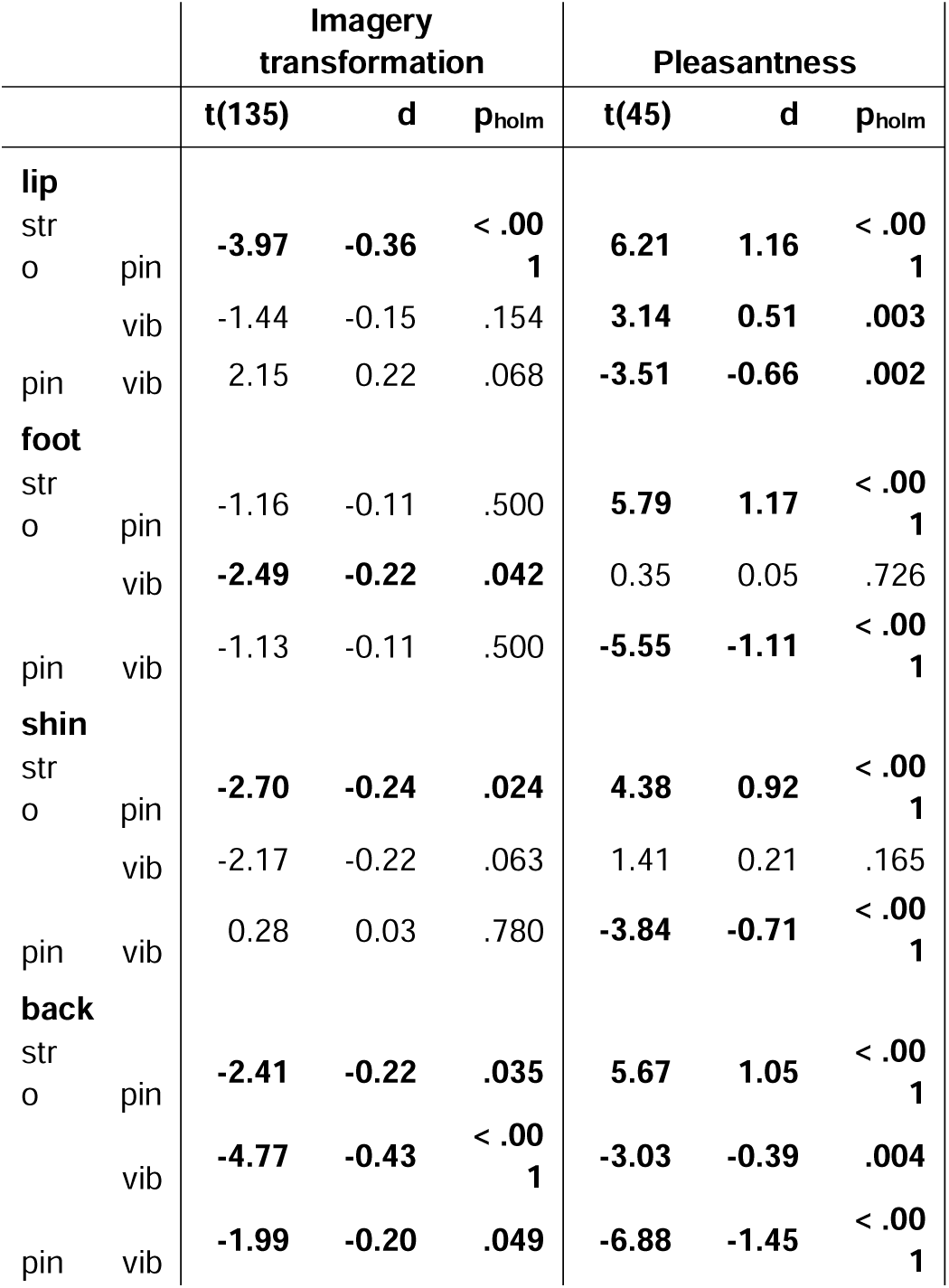
Post-hoc pairwise comparisons for BODY SITE x SENSATION interaction effects for passive tactile imagery transformation and pleasantness. Conditional on factor BODY SITE. d = Cohen’s d; p_holm_ = Holm-Bonferroni corrected p-value adjusted for comparing a family of 3; stro = stroking; pin = pinching; vib = vibration. Significant differences are printed in bold bolded.

Additional conditional comparisons based on factor SENSATION allowed us to compare imagery ability between body sites for each of the sensations (Fig.3H., statistics in Suppl. Table 4). Interestingly, there were no significant differences between body sites for transformation of vibration. Transformation of pinching was significantly higher for the lip than all other body sites. For stroking however, transformation of the lip was only significantly higher than the back. Moreover, transformation of stroking was significantly higher for the foot than the back and the shin.

### Pleasantness of tactile imagery

We additionally explored how pleasantness varies across different types of imagined touch. Descriptives and statistics for overall mean pleasantness scores for active and passive tactile imagery are listed in Table 1. When taking all items together, active tactile imagery was rated as significantly more pleasant than passive tactile imagery, and both were rated above neutral. There was no significant correlation between active and passive tactile imagery pleasantness.

#### Active tactile imagery

For active tactile imagery, pleasantness did not vary significantly across OBJECTS (F(2,90) = 1.70, p = .189, η_p_^2^ = .04, Fig.4A.), PROPERTIES (F(3,135) = 1.66, p = .179, η_p_^2^ = .04, Fig.4B.) or their interaction (F(6,250) = 1.81, p = .104, η_p_^2^ = .04, H-F corrected). This indicates that experienced pleasantness was relatively stable across different types of active touch.

**Figure 4.**
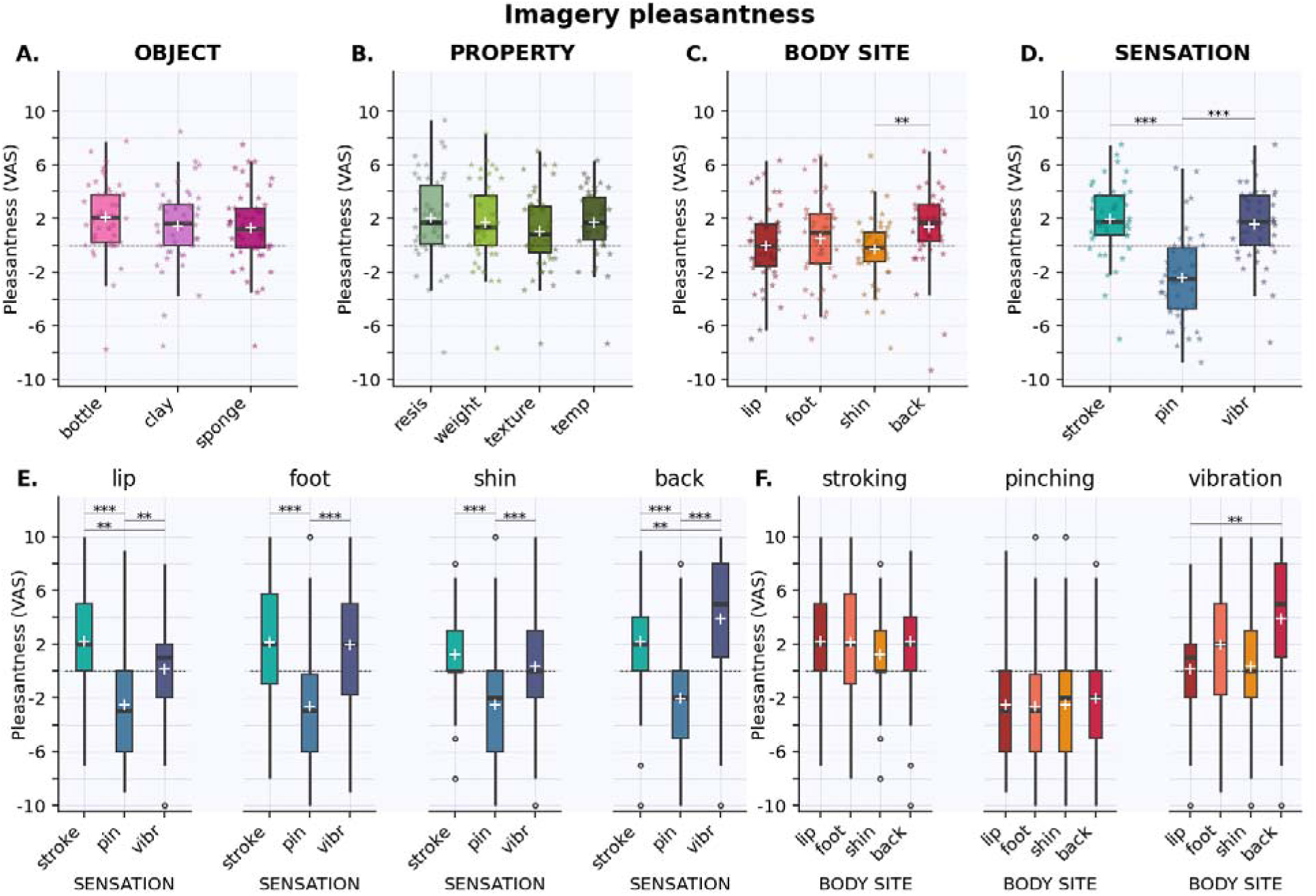
PLEASANTNESS of active (A.,B.) and passive (C.-F.) tactile imagery. Plots show main effects of OBJECT and PROPERTY (A.B., both non-significant) and significant effects of BODY SITE (C.) and SENSATION (D.). Significant interaction effect of BODY SITE x SENSATION is visualized per body site (E.) and per sensation (F.). Pleasantness scores range from -10 (unpleasant) to 10 (pleasant). The dashed line in each plot denotes the neutral midpoint (0). Boxes represent the interquartile ranges (IQR (from the 25^th^ (Q1) to the 75^th^ (Q3) percentile); whiskers show the minimum within Q1 -1.5 times the IQR and maximum within Q3 + 1.5 times the IQR. The black lines show the median, and the white + the mean per condition. = individual data points, jittered for visibility purposes; ◦ **=** outlier; ** significant difference at p_holm_ ≤.01; *** significant difference at p_holm_ ≤.001. stroke = stroking; pin = pinching; vibr = vibration; VAS = Visual Analogue Scale.

#### Passive tactile imagery

Unlike for active tactile imagery, pleasantness did vary significantly across different types of passive tactile imagery. There was a significant main effect of BODY SITE (F(2,112) = 4.92, p = .005, η_p_^2^ = .10, H-F corrected, Fig.4C.). Post-hoc pairwise comparisons revealed that overall passive imagery was significantly more pleasant for the back than the shin (t(44) = 3.95, p_holm_ = .002, d = 0.41). None of the other body sites’ pleasantness differed significantly from each other (Suppl. Table 7).

Pleasantness also differed significantly across passive tactile sensations (F(2,69) = 43.95, p < .001, η_p_^2^ = .50, H-F corrected, Fig.4D.). Post-hoc tests revealed that, as expected, pinching was significantly less pleasant than stroking (t(44) = 6.95, p_holm_ < .001, d = 1.10) and vibration (t(44) = 7.56, p_holm_ < .001, d = 0.98). Stroking and vibration did not differ significantly (t(44) = 1.08, p_holm_ = .285, d = 0.10).

Finally, BODY SITE and SENSATION interacted significantly (F(6,270) = 4.33, p < .001, η_p_^2^ = .09), indicating that the pleasantness of sensations depended on the specific body site. We first conducted conditional comparisons based on factor BODY SITE to compare sensations within body sites (Fig.4F., statistics in Table 3). As expected, pinching was significantly less pleasant than stroking and vibration for all body sites. Different patterns emerged when comparing vibration and stroking. Whereas stroking was significantly *more* pleasant than vibration for the lip, the back showed a reversed pattern, with stroking being *less* pleasant than vibration. Pleasantness of stroking and vibration did not differ significantly for foot and shin.

We additionally performed conditional comparisons based on factor SENSATION to allow comparison of body sites for each of the sensations (Fig.4E., statistics in Suppl.Table.6). Interestingly, pleasantness did not differ between body sites for stroking and pinching. For vibration however, pleasantness was significantly higher for the back than the lip and the shin, and showed a trend in the same direction compared to the foot.

### Correlation between tactile imagery ability and pleasantness

We explored whether tactile imagery ability is related to imagery pleasantness, we correlated imagery ability and pleasantness with each other.

#### Active tactile imagery

For active tactile imagery, overall pleasantness showed significant positive correlations with overall vividness, maintenance and transformation, with medium effect sizes (see Fig.5A.-C. for statistics). Individuals for whom overall pleasantness of active tactile imagery was higher, also had more vivid imagery, with higher transformation and maintenance.

**Figure 5.**
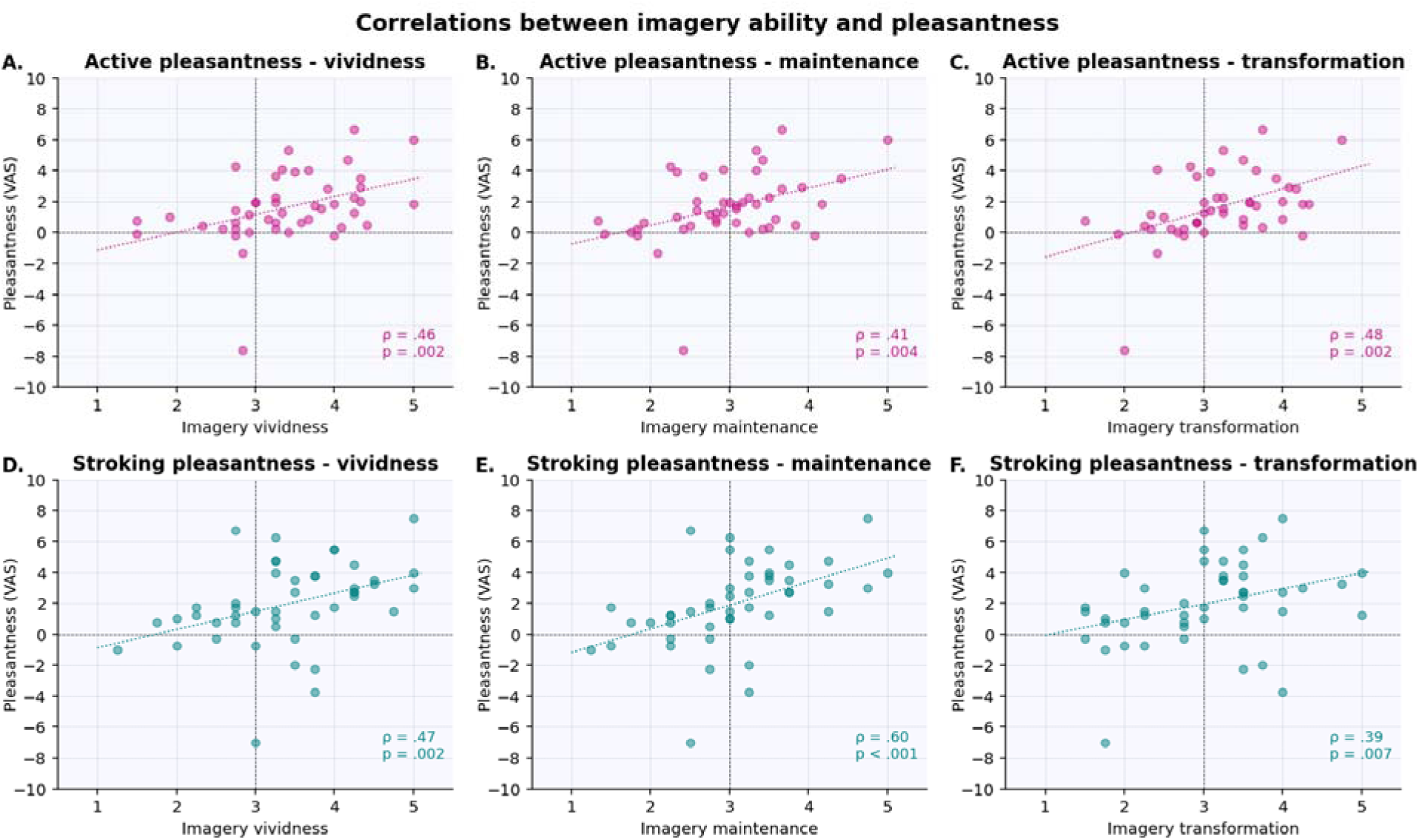
Correlations between imagery ability scores and pleasantness. Overall mean pleasantness of active tactile imagery is significantly positively correlated with overall mean active tactile imagery vividness (A.), maintenance (B.) and transformation (C.). For passive tactile imagery, pleasantness of stroking is similarly significantly and positively correlated with vividness (D.), maintenance (E.) and transformation (F.) of stroking (collapsed across body sites). Pleasantness scores range from -10 (unpleasant) to 10 (pleasant), imagery ability scores range from 1 to 5, with higher scores reflecting higher imagery ability. The black dashed cross in the center shows the neutral midpoint (0) for pleasantness and the imagery ability scale midpoint (3). Dashed colored lines depict trendlines. p = p_holm_, Holm-Bonferroni corrected p-value (n=3); ρ = Spearman’s rho; r = Pearson’s r; VAS = Visual Analogue Scale.

#### Passive tactile imagery

Since pleasantness varied significantly across passive sensations (with both negative (unpleasant) and positive (pleasant) scores), we explored the relationship between imagery ability and pleasantness per sensation. For stroking, pleasantness correlated significantly with imagery vividness, maintenance and transformation, with medium to large effect sizes (see Fig.5D.-F. for statistics). Individuals who rated stroking as more pleasant also reported higher imagery vividness, maintenance and transformation for stroking. Interestingly pleasantness of pinching and vibration did not correlate significantly with any imagery processing component (Suppl.Table.8).

## DISCUSSION

This study aimed to provide a comprehensive assessment of tactile imagery covering the full spectrum of our multidimensional sense of touch. This study was based on O’Dowd and colleagues’ (2022) earlier work, extending it with additional tactile sensations (temperature, pinching, vibration) and imagery processing components (i.e. imagery maintenance and transformation rather than vividness alone). We asked participants to imagine active and passive touch. For active touch, participants imagined feeling various properties (the resistance, temperature, texture and weight) of three different objects, while for passive touch they imagined stroking, pinching and vibration across different body sites. A subset of participants additionally rated experienced pleasantness for each imagined sensation. Our results show that imagery ability and pleasantness vary across different active and passive tactile sensations, and across individuals.

The majority of participants showed at least moderate imagery ability for both active and passive touch, indicating that people are generally able to imagine different types of touch. Nevertheless, individual variation was substantial as scores were found across both ends of the scales, confirming previous reports (O’ Dowd et al., 2022). Whereas O’Dowd (2022) reported that imagery of active and passive touch were only moderately correlated, we found strong correlations between active and passive touch across imagery processing components. The higher correlation between active and passive tactile imagery in the current study may stem from design differences: whereas O’Dowd used vividness for active, and sensitivity for passive tactile imagery, here we used the same imagery scales (i.e. vividness, maintenance, transformation) for both.

Active imagery ratings varied significantly across object-property combinations, with certain properties eliciting stronger imagery for one object, but weaker imagery for another (and vice versa). Our use of geometrically distinct objects, and the addition of temperature yielded important new insights. Whereas O’Dowd and colleagues (2022) reported texture and surface compliance (analogous to our resistance) as the most vivid properties, we found relatively weak imagery for resistance (except for clay), alongside markedly different patterns across objects. Although we replicated O’Dowd’s relatively *strong* imagery for texture of the sponge, texture elicited relatively *weak* imagery for the bottle. Moreover, whereas temperature and weight were relatively weak for sponge and clay, they were stronger for the bottle. These findings indicate that it is easier to imagine touching certain features than others, and that this is object-specific. This aligns with research on tactile object perception, which suggests that during haptic exploration, an object’s material, geometry and contact conditions jointly shape the salience of its individual properties (Bergmann Tiest, 2010). More salient properties in turn become part of lasting object representations (Klatzky et al., 1987). Our results indicate that this object-specific feature salience translates to tactile imagery. This highlights the importance of including a variety of objects in paradigms assessing active tactile imagery.

Passive touch imagery ratings differed significantly across body sites and sensations. Imagery ability was generally higher for body sites with higher tactile sensitivity. Though we used slightly different outcome measures than O’Dowd (2022) (assessing imagery processing components rather than sensitivity), we did replicate their finding of higher ratings for imagined stroking at more sensitive body sites. Imagery of pinching and vibration generally followed a similar pattern, with higher imagery ratings for body sites with higher tactile sensitivity. The one exception was transformation of vibration, for which body sites did not differ significantly. This may be due to slightly different transformation instructions for the lip compared to the other body sites (i.e. from constant to pulsating vibration, vs. an increase in pressure, respectively). Another contributing factor may be variation in familiarity with the vibration sensations, depending on experience with an electric toothbrush (for the lip) or massage gun (for foot, shin and back). Familiarity with a sensation influences imagery vividness in other sensory modalities, where higher exposure to, or familiarity with, a specific sensation is associated with more vivid/accurate imagery of that sensation (Croijmans et al., 2020; Guillot & Collet, 2005; Lacey & Sathian, 2013). This explanation is supported by the fact that numerous participants reported (unprompted) that it was harder to imagine unfamiliar sensations. These factors may have reduced the lip’s ‘tactile sensitivity advantage’ seen in stroking and pinching. Interestingly, stroking generally elicited significantly less vivid imagery and was harder to transform than the other passive sensations, though their maintenance did not differ. This may be due to several factors. First, familiarity with the stimulus may similarly be involved: stroking with cotton wool is less commonly encountered than pinching between fingernails and vibration of a massage gun or electric toothbrush. Second, pinching and vibration are more salient sensations than stroking. Especially pinching is close to an acute pain stimulus, which is known to attract attention (Eccleston & Crombez, 1999). Finally, stroking is dynamic (spatially), where pinching and vibration are spatially static. These involve different somatosensory processes (Timmermann et al., 2000) and it may be more difficult to generate, inspect and transform a dynamic tactile sensation than a static one. When looking at our active touch findings, properties that require static contact (weight/temperature), however, did not show consistently stronger imagery compared to properties that require dynamic exploration in the form of lateral motion (texture) and pressure (resistance) (Lederman & Klatzky, 1987). One possible contributing factor for this discrepancy between active and passive tactile imagery may be that during active haptic exploration of an object, the specific tactile input received by the fingers changes as they run along the object, but the spatial location of that tactile input stays mostly the same (e.g. the fingertips). In contrast, during passive stroking, the type (i.e. pressure, material, velocity, etc.) of tactile input stays constant, but the spatial displacement along the skin is relatively greater. Taken together, these variations in imagery strength across body sites and sensations highlight the complexity of passive tactile imagery, further illustrating the need for inclusion of a variety of sensations and body sites when assessing tactile imagery ability.

An important novelty in the current study was that we included the imagery processing components maintenance and transformation, rather than focusing solely on imagery vividness. Overall, we observed fairly similar patterns for the three imagery components. Correlations between components were strong for both active and passive tactile imagery: people with more vivid imagery also reported higher maintenance and transformation, and people with higher maintenance had higher transformation. Though classically the components are considered to be separate aspects of imagery ability (Kosslyn, 1995; Kosslyn et al., 1984), empirical studies have shown that dissociations between them vary (Arnold et al., 2025; De Beni et al., 2007; Moreno-Verdú et al., 2025). While the current study suggests closer relations between them, the correlations are not 1-to-1, and components show partly different patterns across different types of active and passive touch. For example, transformation appears to be more sensitive to differences in passive touch across body sites and tactile sensations. It confirms that although related, the different imagery processing components cannot be equated, indicating that assessment of imagery ability should not be restricted to vividness alone, as relevant nuances may be lost.

In a subset of participants, we also assessed hedonic experience. Overall, active tactile imagery was rated significantly more pleasant than passive tactile imagery. This is likely because we did not include universally unpleasant active tactile sensations – as reflected in constant pleasantness across objects and properties – whereas passive touch included a painful sensation (i.e. pinching). which is generally experienced as unpleasant. Imagined pinching was indeed experienced as significantly less pleasant than stroking and vibration across body sites. Based on affective touch literature (McGlone et al., 2014), we expected stroking to be the most pleasant passive sensation. Experienced pleasantness of stroking and vibration however only differed significantly for the lip and the lower back, with opposite patterns. Whereas stroking was *more* pleasant than vibration for the lip, it was *less* pleasant than vibration for the lower back. This is likely due to positive associations with the massage gun to the back. Moreover, our stroking material was cotton wool, a material that people are likely less familiar with, and which differs from materials typically used in affective touch research (i.e. soft brushes or hands (Taneja et al., 2021)). More familiar and affective materials may therefore increase experienced pleasantness of imagined stroking, as seen in previous studies where imagined stroking was more pleasant when it was affective, in line with tactile stimulation (Lustenhouwer & Meijer, 2026; Tagini et al., 2023). Interestingly, imagery ability and pleasantness were moderately to strongly related. Individuals with higher overall vividness, maintenance and transformation for active tactile imagery generally experienced it as more pleasant. For passive tactile imagery, ability and pleasantness only correlated for stroking: individuals with higher imagery processing component scores for being stroked reported higher pleasantness scores for stroking. Interestingly, individuals with greater exposure to affective touch find this type of touch more pleasant during tactile stimulation (Sailer & Ackerley, 2019). This link between exposure and hedonic experience of affective touch and the proposed link between familiarity and imagery strength (Croijmans et al., 2020; Guillot & Collet, 2005; Lacey & Sathian, 2013), may mediate the correlation between imagery ability and pleasantness we find here. It is difficult to determine whether imagery strength drives experienced pleasantness, or whether more pleasant tactile sensations elicit stronger imagery. Regardless of directionality, this links warrants further investigation, especially considering current interest in clinical applications for affective tactile imagery, where pleasantness is an important factor (Lustenhouwer & Meijer, 2026; Tagini et al., 2023).

This study has several strengths and limitations. Focusing on our design first: we provide a comprehensive assessment of tactile imagery that reflects the multidimensionality of our sense of touch. Our within-subject design allowed for direct comparison across tactile sensations. Trained assessors instructed participants and recorded data in a controlled setting, minimizing noise and missing data. A design-related limitation is the use of self-reported rating scales, which may be subject to response bias (Allbutt et al., 2011). In absence of objective imagery ability measures that are cost-effective and easily applicable across research and clinical settings, there is a need for modality-specific, theoretically driven questionnaires that include vividness as well as process-oriented items (Lacey & Lawson, 2013). Here we aimed to work towards just that, starting from tactile literature, and assessing core imagery processing components beyond vividness. Because of this, we did not randomize or counterbalance items, which may have introduced fatigue, diminished attention, and learning effects. To minimize effects of fatigue and diminished attention, participants were encouraged to take self-paced breaks when needed. Second, related to our sample: we assessed tactile imagery in a large, but homogenous healthy sample (n=136), with an overrepresentation of young, female psychology students in the Netherlands. Although there are currently no studies on the influence of gender, age, or level of educational on *tactile* imagery ability, studies in other sensory modalities report variation in ability within and between modalities related to these factors, though results are somewhat inconsistent (Campos, 2014; Conson et al., 2020; Fierro-Marrero et al., 2025; Floridou et al., 2022). Given these findings in other imagery modalities, gender and age effects in tactile processing (Da Silva et al., 2014; Godde et al., 2018; Neely & Burström, 2006; Powell et al., 2025; Sehlstedt et al., 2016; Tirrell et al., 2025), and cross-cultural differences in stimulus familiarity, generalizability of our results may be limited. Likewise, our pleasantness data cover a smaller sample (n=46), which means our pleasantness analyses may be underpowered. Larger, more diverse samples are needed to replicate and expand our findings on both imagery ability and hedonic experience. Finally, we aimed to parse out imagery ability specific to the tactile modality, which is why we instructed participants to focus on the tactile sensation and refrain from using visual information during their imagery. Despite these instructions, we could not objectively monitor visual imagery use and its interference with tactile imagery. In fact, many participants reported difficulties with blocking visual imagery during their tactile imagery. This may have influenced our results, though it is difficult to say in what way. Research in other modalities has reported enhancement of imagery when multiple modalities are combined (Holmes & Collins, 2001). This is of course the case for motor imagery of the exploratory procedures, such as lateral motion, pressure, unsupported holding etc., as well. As these involve active hand and finger movements, it is inevitable that active tactile imagery also contains a motor imagery component, particularly when it comes to imagery of surface features (Klatzky et al., 1991; O’ Dowd et al., 2022).

Taken together, our results show that imagery ability varies significantly across different tactile sensations, both active and passive. Most existing (multisensory) imagery ability questionnaires (e.g. Bett’s Questionnaire Upon Mental Imagery, Plymouth Sensory Imagery Questionnaire) include only a few items on vividness of touch and give rather vague instructions that are not focused on specific features of a tactile sensation (Andrade et al., 2014; Sheehan, 1967). Our findings indicate that these limited assessments are likely to give an incomplete picture of tactile imagery ability. Given the recently increased interest in tactile imagery, there is a need for a comprehensive tool that does justice to the breadth and multidimensional nature of our sense of touch. The 72-item assessment we used here provides a first step towards a modality-specific tactile imagery ability assessment tool that can be applied in research and clinical settings alike. To enable uniform assessment of tactile imagery ability that truly captures individual variability rather than other factors, future studies should focus on the following issues. First, the familiarity of different tactile sensations should be mapped to inform selection of tactile sensations with relatively stable familiarity (either familiar or unfamiliar) across individuals, to ensure that we are capturing variations in tactile imagery ability rather than variations in familiarity with specific sensations. Second, the link between hedonic experience and imagery ability should be investigated further, so that it can inform item selection, to include both pleasant and unpleasant tactile sensations. These factors should ideally be studied across different populations (healthy, patient, cross-cultural). Finally, the relation between imagery in the tactile domain and other sensory modalities should be explored further, with particular interest in visual imagery and motor imagery.

To conclude, our results confirm that individuals are generally able to imagine active and passive tactile sensations, with considerable individual variability. Importantly, imagery ability varies across objects and their properties, and across sensations and body sites, with a general advantage for body sites with higher tactile sensitivity. Performance on the maintenance and transformation components of tactile imagery were highly related to vividness, but not quite identical, suggesting that it is relevant to include all three imagery processing components when assessing tactile imagery. Pleasantness was probed in a subsample and correlated with imagery ability, an observation that requires further exploration in future studies. Overall, this study has provided important additional information about the multidimensionality of tactile imagery and we aim to use this as a first step to develop a comprehensive assessment tool of this vastly understudied imagery domain.

## DATA STATEMENT

All data and analyses files (Python and JASP files) are available publicly available at https://doi.org/10.24416/UU01-CVUXP1

## CREDIT AUTHORSHIP CONTRIBUTION STATEMENT

Renee Lustenhouwer: Writing – review & editing, Writing – original draft, Visualization, Supervision, Software, Resources, Project administration, Methodology, Investigation, Formal analysis, Data curation, Conceptualization. H Chris Dijkerman: Writing – review & editing, Writing – original draft, Methodology, Conceptualization.

## ACKNOWLEDGMENTS

We thank Christian Cano, Fotini Eracleous, Kaja Kleszczewska, Davide Neagoe, Costantino Rossotto, and Mert Saraçoğlu for their role in data collection. We also would like to thank Fiona Newell and Alan O’Dowd for their input during the conceptualization of the study and the useful discussions concerning the interpretation of the results.

