## Supplementary Materials for "Capturing the imagination: mapping imagery ability across our multidimensional sense of touch"

### Tactile Imagery Ability Assessment Task

#### Section 1: Active tactile imagery

Instructions:

Please imagine the sensation of touching the following objects. Rate the vividness of the imagined sensation on a scale from 1 (*No imagery [I only know that I am thinking about the sensation])* to 5 (*Very detailed and vivid [the imagery of the sensation is as vivid as a real stimulus]*). Additionally, you will be asked to rate the controllability of the imagined sensation on a scale from 1 (*Unable to control the imagery of the sensation [I cannot manipulate or change the imagery of the sensation at all])* to 5 (*Completely able to control the imagery of the sensation [I can fully manipulate or change the imagery of the sensation as intended]*) and the maintenance of the imagined sensation on a scale from 1 (*Unable to maintain the imagery of the sensation [the imagery of the sensation fades quickly and cannot be sustained]*) to 5 (*Completely able to maintain the imagery of the sensation [the imagery of sensation is steady and does not fade over time]*). Please focus solely on the feeling of touch, make sure to NOT use any visual information.

##### Vividness scale

| 1 | 2 | 3 | 4 | 5 |
|---|---|---|---|---|
| No imagery (I only know that I am thinking about the sensation) | Vague and unclear (the imagery of the sensation is faint and lacks detail) | Relatively clear and vivid (the imagery of the sensation has some detail but is not fully vivid) | Clear and vivid (the imagery of the sensation is detailed and realistic) | Very detailed and vivid (the imagery of the sensation is as vivid as a real stimulus) |

##### Transformation scale

| 1 | 2 | 3 | 4 | 5 |
|---|---|---|---|---|
| Unable to control the imagery of the sensation (I cannot manipulate or change the imagery of the sensation at all) | Slightly able to control the imagery of the sensation (I can make minimal changes to the imagery of the sensation) | Moderately able to control the imagery of the sensation (I can somewhat manipulate or change the imagery of the sensation) | Very able to control the imagery of the sensation (I can easily manipulate or change the imagery of the sensation) | Completely able to control the imagery of the sensation (I can fully manipulate or change the imagery of the sensation as intended) |

##### Maintenance scale

| 1 | 2 | 3 | 4 | 5 |
|---|---|---|---|---|
| Unable to maintain the imagery of the sensation (the imagery of the sensation fades quickly and cannot be sustained) | Slightly able to maintain the imagery of the sensation (the imagery of the sensation fades but can be briefly sustained) | Moderately able to maintain the imagery of the sensation (the imagery of the sensation can be sustained for a short period with some effort) | Very able to maintain the imagery of the sensation (The imagery of the sensation can be sustained for a longer period with minimal fading) | Completely able to maintain the imagery of the sensation (the imagery of sensation is steady and does not fade over time) |

Supplementary materials to *Capturing the imagination: mapping imagery ability across our multidimensional sense of touch* – Lustenhouwer & Dijkerman

1. Plastic Bottle

1.1. Close your eyes. Imagine holding a plastic bottle filled with cold water in your hands. How vividly can you imagine its cold surface?

1.2. Close your eyes. Now imagine again the sensation of holding the plastic bottle filled with cold water in your hands and hold this sensation for 5 seconds. How well are you able to maintain the sensation of its cold surface?

1.3. Close your eyes. Now imagine that the water bottle is gradually getting hotter as you are holding it. How well are you able to control the change of this sensation?

2.1 Close your eyes. Imagine squeezing an empty plastic bottle in your hand. How vividly can you imagine the resistance (softness or hardness of the surface) it gives as you squeeze?

2.2 Close your eyes. Now imagine again the sensation of squeezing the empty bottle in your hand and hold this sensation for 5 seconds. How well are you able to maintain the sensation of squeezing the water bottle?

2.3 Close your eyes. Now imagine that the water bottle is gradually being filled with water and therefore making it harder to squeeze. How well are you able to control the change of this sensation?

3.1 Close your eyes. Imagine the smooth texture of an empty plastic bottle as your fingers repeatedly run along its surface. How vividly can you imagine the sensation of its texture?

3.2 Close your eyes. Now imagine again the smooth texture of the plastic bottle and hold this sensation for 5 seconds. How well are you able to maintain the sensation of the bottle's texture?

3.3 Close your eyes. Now imagine slowly squeezing the bottle, gradually making its surface less smooth and more wrinkly. How well are you able to control the change of this sensation?

4.1 Close your eyes. Imagine the strength needed to lift an empty water bottle. How vividly can you imagine the sensation of its weight?

4.2 Close your eyes. Now imagine again the strength needed to lift an empty water bottle and hold this sensation for 5 seconds. How well are you able to maintain the sensation of the bottle's weight?

4.3 Close your eyes. Now imagine the bottle is gradually getting heavier as it fills up with water. How well are you able to control the change of this sensation?

2. Modelling Clay

1.1 Close your eyes. Imagine touching modelling clay that feels cold to the touch. How vividly can you imagine the sensation of its temperature?

1.2 Close your eyes. Now imagine again touching the clay that feels cold to the touch and hold this sensation for 5 seconds. How well are you able to maintain the sensation of the clay's temperature?

1.3 Close your eyes. Now imagine the clay gradually getting warmer to the touch. How well are you able to control the change of this sensation?

2.1 Close your eyes. Imagine squeezing the modelling clay with your hands. How vividly can you imagine the resistance (softness or hardness of the surface) it gives as you squeeze?

2.2 Close your eyes. Now imagine again squeezing the modelling clay with your hands and hold this sensation for 5 seconds. How well are you able to maintain the sensation of the clay's resistance?
2.3 Close your eyes. Now imagine the clay gradually getting softer to the touch as more water is added to it. How well are you able to control the change of this sensation?

3.1 Close your eyes. Imagine the texture of modelling clay. How vividly can you imagine the sensation of its texture?
3.2 Close your eyes. Now imagine again the texture of modelling clay and hold this sensation for 5 seconds. How well are you able to maintain the sensation of the clay's texture?
3.3 Close your eyes. Now imagine the sensation of the texture of the clay as it gradually dries out making its surface rougher and 'chalky'. How well are you able to control the change of this sensation?

4.1 Close your eyes. Imagine the strength needed to lift a clump of clay. How vividly can you imagine the sensation of its weight?
4.2 Close your eyes. Now imagine the strength needed to lift the clump of clay and hold this sensation for 5 seconds. How well are you able to maintain the sensation of the clay's weight?
4.3 Close your eyes. Now imagine the weight of the clay gradually getting heavier as more clay is added to the clump you are holding. How well are you able to control the change of this sensation?

#### 3. Sponge

1.1 Close your eyes. Imagine holding a wet sponge soaked in cold water. How vividly can you imagine its temperature?
1.2 Close your eyes. Now imagine again holding the wet sponge soaked in cold water and hold this sensation for 5 seconds. How well are you able to maintain the sensation of its cold surface?
1.3 Close your eyes. Now imagine the sponge gradually warming up in your hands. How well are you able to control the change of this sensation?

2.1 Close your eyes. Imagine pressing against a dry sponge and feeling its resistance (softness or hardness of surface). How vividly can you imagine the resistance as you press against the sponge?
2.2 Close your eyes. Now imagine again pressing against a dry sponge that is harder to press and hold this sensation for 5 seconds. How well are you able to maintain the sensation of feeling its resistance?
2.3 Close your eyes. Now imagine that the sponge you are holding is gradually being soaked with water, making it softer to press. How well are you able to control the change of this sensation?

3.1 Close your eyes. Imagine the texture of the soft side of the sponge. How vividly can you imagine the sensation of its texture?
3.2 Close your eyes. Now again imagine the texture of the soft side of the sponge and hold this sensation for 5 seconds. How well are you able to maintain the sensation of its texture?
3.3 Close your eyes. Now imagine running your finger from the soft side of the sponge towards the rough side of the sponge. How well are you able to control the change of this sensation?

4.1 Close your eyes. Imagine the strength needed to lift a dry sponge. How vividly can you imagine the sensation of its weight?
4.2 Close your eyes. Now imagine again the strength needed to lift the dry sponge and hold this sensation for 5 seconds. How well are you able to maintain the sensation of its weight?
4.3 Close your eyes. Now imagine the sponge becoming heavier as it soaks in water. How well are you able to control the change of this sensation?

### Section 2: Passive tactile imagery

<u>Instructions:</u>

Please imagine the following sensations on the specified body parts. Rate the vividness of the imagined sensation on a scale from 1 (*No imagery [I only know that I am thinking about the sensation])* to 5 (*Very detailed and vivid [the imagery of the sensation is as vivid as a real stimulus]*). Additionally, you will be asked to rate the controllability of the imagined sensation on a scale from 1 (*Unable to control the imagery of the sensation [I cannot manipulate or change the imagery of the sensation at all])* to 5 (*Completely able to control the imagery of the sensation [I can fully manipulate or change the imagery of the sensation as intended]*) and the maintenance of the imagined sensation on a scale from 1 (*Unable to maintain the imagery of the sensation [the imagery of the sensation fades quickly and cannot be sustained]*) to 5 (*Completely able to maintain the imagery of the sensation [the imagery of sensation is steady and does not fade over time]*). Please focus solely on the feeling of touch, make sure to NOT use any visual information.

**Vividness scale**

| 1 | 2 | 3 | 4 | 5 |
|---|---|---|---|---|
| No imagery (I only know that I am thinking about the sensation) | Vague and unclear (the imagery of the sensation is faint and lacks detail) | Relatively clear and vivid (the imagery of the sensation has some detail but is not fully vivid) | Clear and vivid (the imagery of the sensation is detailed and realistic) | Very detailed and vivid (the imagery of the sensation is as vivid as a real stimulus) |

**Transformation scale**

| 1 | 2 | 3 | 4 | 5 |
|---|---|---|---|---|
| Unable to control the imagery of the sensation (I cannot manipulate or change the imagery of the sensation at all) | Slightly able to control the imagery of the sensation (I can make minimal changes to the imagery of the sensation) | Moderately able to control the imagery of the sensation (I can somewhat manipulate or change the imagery of the sensation) | Very able to control the imagery of the sensation (I can easily manipulate or change the imagery of the sensation) | Completely able to control the imagery of the sensation (I can fully manipulate or change the imagery of the sensation as intended) |

#### Maintenance scale

| 1 | 2 | 3 | 4 | 5 |
|---|---|---|---|---|
| Unable to maintain the imagery of the sensation (the imagery of the sensation fades quickly and cannot be sustained) | Slightly able to maintain the imagery of the sensation (the imagery of the sensation fades but can be briefly sustained) | Moderately able to maintain the imagery of the sensation (the imagery of the sensation can be sustained for a short period with some effort) | Very able to maintain the imagery of the sensation (The imagery of the sensation can be sustained for a longer period with minimal fading) | Completely able to maintain the imagery of the sensation (the imagery of sensation is steady and does not fade over time) |

##### 1. Lower Lip

1.1 Close your eyes. Imagine the feeling of cotton wool gently (and lightly) stroking your lower lip. The stroking moves from side to side within 1 second. How vividly can you imagine this sensation?

1.2 Close your eyes. Imagine that the gentle stroking on your lower lip smoothly and gradually increases in pressure. The direction and speed of the strokes stay the same. How well are you able to control the change of this sensation?

1.3 Close your eyes. Imagine again the feeling of cotton wool gently (and lightly) stroking your lower lip. The stroking moves from side to side within 1 second. Now hold this stroking sensation for 5 seconds. How well are you able to maintain this sensation?

1.4 Close your eyes. Imagine the feeling of someone lightly pinching your lower lip with their nails, creating a slight feeling of pain. How vividly can you imagine this sensation?

1.5 Close your eyes. Imagine that the light pinching on your lower lip gradually increases in pressure, making it more painful. How well are you able to control the change of this sensation?

1.6 Close your eyes. Imagine again the feeling of someone lightly pinching your lower lip with their nails, creating a slight feeling of pain. Now hold this pinching sensation for 5 seconds. How well are you able to maintain this sensation?

1.7 Close your eyes. Imagine the feeling of an electrical toothbrush lightly and continuously vibrating on your lower lip. How vividly can you imagine this sensation?

1.8 Close your eyes. Imagine again the light continuous vibrating sensation of the toothbrush on your lower lip. Now it changes into a pulsating vibration. How well are you able to control the change of this sensation?

1.9 Close your eyes. Imagine the feeling of an electrical toothbrush lightly and continuously vibrating on your lower lip. Now hold this sensation for 5 seconds. How well are you able to maintain this sensation?

##### 2 Lower Back

2.1 Close your eyes. Imagine the feeling of cotton wool gently (and lightly) stroking your lower back. The strokes move from left to right on your lower back area within 1 second. How vividly can you imagine this sensation?

2.2 Close your eyes. Imagine that the gentle stroking on your lower back smoothly and gradually increases in pressure. The direction and speed of the strokes stay the same. How well are you able to control the change of this sensation?

2.3 Close your eyes. Imagine again the feeling of cotton wool gently (and lightly) stroking your lower back. The strokes move from left to right on your lower back area within 1 second. Now hold this stroking sensation for 5 seconds. How well are you able to maintain this sensation?

2.4 Close your eyes. Imagine the feeling of someone lightly pinching your lower back with their nails, creating a slight feeling of pain. How vividly can you imagine this sensation?
2.5 Close your eyes. Imagine that the light pinching on your lower back gradually increases in pressure, making it more painful. How well are you able to control the change of this sensation?
2.6 Close your eyes. Imagine again the feeling of someone lightly pinching your lower back with their nails, creating a slight feeling of pain. Now hold this pinching sensation for 5 seconds. How well are you able to maintain this sensation?

2.7 Close your eyes. Imagine the feeling of a massage gun lightly vibrating on your lower back. How vividly can you imagine this sensation?
2.8 Close your eyes. Imagine that the light vibration of the massage gun on your lower back gradually increases in pressure, making the vibration harder. How well are you able to control the change of this sensation?
2.9 Close your eyes. Imagine again the feeling of a massage gun lightly vibrating on your lower back. Now hold this sensation vibrating sensation for 5 seconds. How well are you able to maintain this sensation?

3 Left Shin

3.1 Close your eyes. Imagine the feeling of cotton wool gently (and lightly) stroking your left shin. The strokes move from just below the knee to just above your ankle within 1 second. How vividly can you imagine this sensation?
3.2 Close your eyes. Imagine that the gentle stroking on your left shin smoothly and gradually increases in pressure. The direction and speed of the strokes stay the same. How well are you able to control the change of this sensation?
3.3 Close your eyes. Imagine again the feeling of cotton wool gently (and lightly) stroking your left shin. The strokes move from just below the knee to just above your ankle within 1 second. Now hold this stroking sensation for 5 seconds. How well are you able to maintain this sensation?

3.4 Close your eyes. Imagine the feeling of someone lightly pinching your left shin with their nails, creating a slight feeling of pain. How vividly can you imagine this sensation?
3.5 Close your eyes. Imagine that the light pinching on your left shin gradually increases in pressure, making it more painful. How well are you able to control the change of this sensation?
3.6 Close your eyes. Imagine again the feeling of someone lightly pinching your left shin with their nails, creating a slight feeling of pain. Now hold this pinching sensation for 5 seconds. How well are you able to maintain this sensation?

3.7 Close your eyes. Imagine the feeling of a massage gun lightly vibrating on your left shin. How vividly can you imagine this sensation?
3.8 Close your eyes. Imagine that the light vibration of the massage gun on your left shin gradually increases in pressure, making the vibration harder. How well are you able to control the change of this sensation?
3.9 Close your eyes. Imagine again the feeling of a massage gun lightly vibrating on your left shin. Now hold this vibrating sensation for 5 seconds. How well are you able to maintain this sensation?

4 Sole of Right Foot

4.1 Close your eyes. Imagine the feeling of cotton wool gently (and lightly) stroking the sole of your right foot. The strokes move from just below the toes in the direction of your heel within 1 second. How vividly can you imagine this sensation?

4.2 Close your eyes. Imagine that the gentle stroking on the sole of your right foot smoothly and gradually increases in pressure. The direction and speed of the strokes stay the same. How well are you able to control the change of this sensation?
4.3 Close your eyes. Imagine the feeling of cotton wool gently (and lightly) stroking the sole of your right foot. The strokes move from just below the toes in the direction of your heel within 1 second. Now hold this stroking sensation for 5 seconds. How well are you able to maintain this sensation?

4.4 Close your eyes. Imagine the feeling of someone lightly pinching the sole of your right foot with their nails, creating a slight feeling of pain. How vividly can you imagine this sensation?
4.5 Close your eyes. Imagine that the light pinching on the sole of your right foot gradually increases in pressure, making it more painful. How well are you able to control the change of this sensation?
4.6 Close your eyes. Imagine again the feeling of someone lightly pinching the sole of your right foot with their nails, creating a slight feeling of pain. Now hold this pinching sensation for 5 seconds. How well are you able to maintain this sensation?

4.7 Close your eyes. Imagine the feeling of a massage gun lightly vibrating on the sole of your right foot. How vividly can you imagine this sensation?
4.8 Close your eyes. Imagine that the light vibration of the massage gun on the sole of your right foot gradually increases in pressure, making the vibration harder. How well are you able to control the change of this sensation?
4.9 Close your eyes. Imagine again the feeling of a massage gun lightly vibrating on the sole of your right foot. Now hold this vibrating sensation for 5 seconds. How well are you able to maintain this sensation?

<u>Additional questions post-questionnaire:</u>

1. Were the tasks easy to understand and easy to execute? Please explain:
2. How easy or difficult was it to use tactile-only imagery without relying on visual information (1=Very easy, 2=Somewhat easy, 3=Neutral, 4=Somewhat difficult, 5=Very difficult)? If you found it difficult, please explain why: ______
3. How often do you engage in the imagination of touch in your daily life (e.g. imagining what it feels like to touch or to be touch by something)? (1=Never, 2=Rarely, 3=Occasionally, 4=Often, 5=Very often)
5. What improvements could be made to better this questionnaire?
6. Were there any other remarkable things you would like to mention?

### Active tactile imagery

#### ***Vividness*** *- additional post-hoc testing*

##### *OBJECT x PROPERTY*

To further explore the significant interaction effect of OBJECT x PROPERTY. We performed additional post-hoc conditional comparisons on factor PROPERTY. This revealed that imagery of the bottle was significantly *less* vivid than the clay and the sponge for resistance, whereas it was significantly *more* vivid than the clay and the sponge for both temperature and weight (Suppl.Table.1, and Suppl.Fig.1). Finally, the sponge was significantly *more* vivid than the bottle and the clay for texture.

*Supplementary Table 1 -* ***Vividness*** *additional post hoc testing for significant OBJECT x PROPERTY interaction effect. Conditional comparisons on factor PROPERTY. Significant differences are printed in bold.* $p_{holm}$ *= Holm-Bonferroni corrected p-value adjusted for comparing a family of 3; SE = standard error; temp = temperature; text = texture; wei = weight.*

| PROPERTY | | | Mean Difference | SE | t(135) | Cohen's d | $p_{holm}$ |
|---|---|---|---|---|---|---|---|
| **resistance** | **Bottle** | **Clay** | **-0.404** | **0.087** | **-4.652** | **-0.376** | **< .001** |
| | | **Sponge** | **-0.301** | **0.086** | **-3.495** | **-0.280** | **0.001** |
| | Clay | Sponge | 0.103 | 0.087 | 1.177 | 0.096 | 0.241 |
| **temp** | **Bottle** | **Clay** | **0.610** | **0.103** | **5.932** | **0.567** | **< .001** |
| | | **Sponge** | **0.574** | **0.083** | **6.889** | **0.533** | **< .001** |
| | Clay | Sponge | -0.037 | 0.111 | -0.332 | -0.034 | 0.740 |
| **text** | **Bottle** | Clay | 0.066 | 0.107 | 0.618 | 0.062 | 0.538 |
| | | **Sponge** | **-0.243** | **0.084** | **-2.873** | **-0.226** | **0.009** |
| | **Clay** | **Sponge** | **-0.309** | **0.098** | **-3.163** | **-0.287** | **0.006** |
| **wei** | **Bottle** | **Clay** | **0.485** | **0.118** | **4.110** | **0.451** | **< .001** |
| | | **Sponge** | **0.382** | **0.089** | **4.316** | **0.355** | **< .001** |
| | Clay | Sponge | -0.103 | 0.124 | -0.830 | -0.096 | 0.408 |

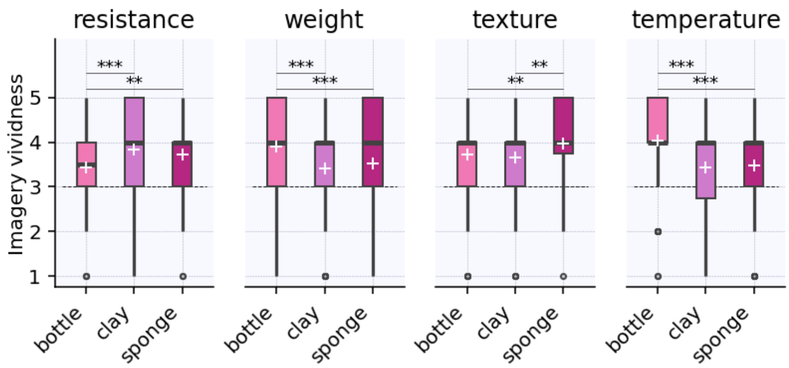


*Supplementary Figure 1 – OBJECT x PROPERTY interaction effect for active tactile imagery* ***vividness.*** *Conditional comparisons on factor PROPERTY. Scores range from 1 to 5, with higher scores denoting higher imagery vividness. Boxes represent the interquartile ranges (IQR (from the 25th (Q1) to the 75th (Q3) percentile); whiskers show the minimum within Q1 -1.5 times the IQR and maximum within Q3 + 1.5 times the IQR. The black lines show the median, and the white + the mean per condition. The dashed line in eachdenotes the scale midpoint (3). ∘ = outlier; **significant difference at* $p_{holm}$ *≤.01; ***significant difference at* $p_{holm}$ *≤.001.*

#### ***Maintenance*** *- additional post-hoc testing*

##### *PROPERTY*

*Supplementary Table 2 - Post Hoc comparisons for significant main effect of PROPERTY for imagery* ***maintenance****. Significant differences are printed in bold.* $p_{holm}$ *= Holm-Bonferroni corrected p-value adjusted for comparing a family of 6; SE = standard error; temp = temperature; text = texture; wei = weight.*

| | | Mean Difference | SE | t (135) | Cohen's d | $p_{holm}$ |
|---|---|---|---|---|---|---|
| **resistance** | temp | -0.174 | 0.071 | -2.438 | -0.148 | 0.080 |
| | **text** | **-0.294** | **0.064** | **-4.619** | **-0.249** | **< .001** |
| | wei | -0.150 | 0.069 | -2.163 | -0.127 | 0.129 |
| temp | text | -0.120 | 0.077 | -1.565 | -0.102 | 0.240 |
| | wei | 0.025 | 0.076 | 0.322 | 0.021 | 0.748 |
| text | wei | 0.145 | 0.074 | 1.956 | 0.123 | 0.158 |

##### *OBJECT x PROPERTY*

To further explore the significant interaction effect of OBJECT x PROPERTY. We performed additional post-hoc conditional comparisons on factor PROPERTY for imagery maintenance. Similar to vividness, the bottle showed significantly *higher* maintenance than clay and sponge for temperature and weight, but significantly *lower* maintenance than the clay for resistance. Unlike vividness, maintenance of resistance was significantly higher for the clay than for the sponge. Imagery maintenance of texture did not differ significantly between the objects (Suppl.Table.3, Suppl.Fig.2).

*Supplementary Table 3 -* ***Maintenance*** *additional post hoc testing for significant OBJECT x PROPERTY interaction effect. Conditional comparisons on factor PROPERTY. Significant differences are printed in bold.* $p_{holm}$ *= Holm-Bonferroni corrected p-value adjusted for comparing a family of 3; SE = standard error; temp = temperature; text = texture; wei = weight.*

| PROPERTY | | | Mean Difference | SE | t(135) | Cohen's d | $p_{holm}$ |
|---|---|---|---|---|---|---|---|
| **resistance** | **Bottle** | **Clay** | **-0.449** | **0.112** | **-4.011** | **-0.380** | **< .001** |
| | | Sponge | -0.162 | 0.107 | -1.511 | -0.137 | 0.133 |
| | **Clay** | **Sponge** | **0.287** | **0.106** | **2.708** | **0.243** | **0.015** |
| **temp** | **Bottle** | **Clay** | **0.618** | **0.104** | **5.916** | **0.524** | **< .001** |
| | | **Sponge** | **0.632** | **0.100** | **6.323** | **0.536** | **< .001** |
| | Clay | Sponge | 0.015 | 0.112 | 0.131 | 0.012 | 0.896 |
| text | Bottle | Clay | -0.088 | 0.108 | -0.815 | -0.075 | 0.832 |
| | | Sponge | -0.147 | 0.095 | -1.551 | -0.125 | 0.370 |
| | Clay | Sponge | -0.059 | 0.106 | -0.553 | -0.050 | 0.832 |
| **wei** | **Bottle** | **Clay** | **0.301** | **0.122** | **2.465** | **0.256** | **0.038** |
| | | **Sponge** | **0.250** | **0.099** | **2.528** | **0.212** | **0.038** |
| | Clay | Sponge | -0.051 | 0.117 | -0.441 | -0.044 | 0.660 |

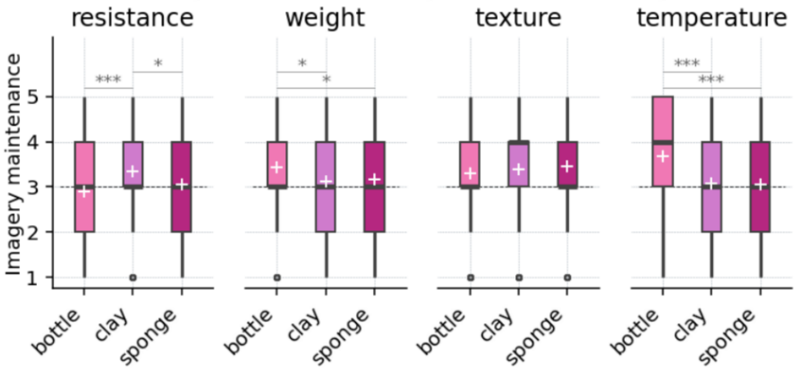


*Supplementary Figure 2 – OBJECT x PROPERTY interaction effect for active tactile imagery* ***maintenance****. Conditional comparisons on factor PROPERTY. Scores range from 1 to 5, with higher scores denoting higher imagery maintenance. Boxes represent the interquartile ranges (IQR (from the 25th (Q1) to the 75th (Q3) percentile); whiskers show the minimum within Q1 -1.5 times the IQR and maximum within Q3 + 1.5 times the IQR. The black lines show the median, and the white + the mean per condition. The dashed line in eachdenotes the scale midpoint (3). ∘ = outlier; *significant difference at* $p_{holm}$ *≤.05; ***significant difference at* $p_{holm}$ *≤.001.*

Supplementary materials to *Capturing the imagination: mapping imagery ability across our multidimensional sense of touch* – Lustenhouwer & Dijkerman

#### ***Transformation** - additional post-hoc testing*

##### *OBJECT*

*Supplementary Table 4 - Post Hoc comparisons for significant main effect of OBJECT for imagery* ***transformation****. Significant differences are printed in bold.* $p_{holm}$ *= Holm-Bonferroni corrected p-value adjusted for comparing a family of 3; SE = standard error.*

| | | Mean Difference | SE | t(134) | Cohen's d | $p_{holm}$ |
|---|---|---|---|---|---|---|
| Bottle | Clay | 0.107 | 0.060 | 1.803 | 0.099 | 0.147 |
| | Sponge | -0.048 | 0.048 | -0.997 | -0.045 | 0.321 |
| **Clay** | **Sponge** | **-0.156** | **0.059** | **-2.657** | **-0.144** | **0.026** |

##### *OBJECT x PROPERTY*

Post-hoc conditional comparisons based on factor PROPERTY revealed that imagery transformation differed between object-property pairs (Suppl.Fig.3, Suppl.Table.5). Transformation of resistance and texture was significantly higher for the sponge than the bottle. The opposite pattern emerged for temperature and weight, where transformation was significantly *lower* for the sponge compared to the bottle. Transformation of the clay was significantly lower than the sponge for texture, and significantly lower than the bottle for weight. For temperature, transformation of the clay showed a trend towards being lower than the bottle.

*Supplementary Table 5 –* ***Transformation*** *additional post hoc testing for significant OBJECT x PROPERTY interaction effect. Conditional comparisons on factor PROPERTY. Significant differences are printed in bold.* $p_{holm}$ *= Holm-Bonferroni corrected p-value adjusted for comparing a family of 3; SE = standard error; temp = temperature; text = texture; wei = weight.*

| PROPERTY | | | Mean Difference | SE | t(134) | Cohen's d | $p_{holm}$ |
|---|---|---|---|---|---|---|---|
| **resistance** | **Bottle** | Clay | -0.126 | 0.113 | -1.115 | -0.116 | 0.368 |
| | | **Sponge** | **-0.281** | **0.104** | **-2.695** | **-0.260** | **0.024** |
| | Clay | Sponge | -0.156 | 0.117 | -1.335 | -0.144 | 0.368 |
| **temp** | **Bottle** | Clay | 0.244 | 0.119 | 2.059 | 0.226 | 0.083 |
| | | **Sponge** | **0.326** | **0.102** | **3.200** | **0.301** | **0.005** |
| | Clay | Sponge | 0.081 | 0.113 | 0.719 | 0.075 | 0.473 |
| **text** | **Bottle** | Clay | -0.133 | 0.117 | -1.140 | -0.123 | 0.256 |
| | | **Sponge** | **-0.637** | **0.102** | **-6.262** | **-0.589** | **< .001** |
| | **Clay** | **Sponge** | **-0.504** | **0.102** | **-4.918** | **-0.466** | **< .001** |
| **wei** | **Bottle** | **Clay** | **0.444** | **0.104** | **4.255** | **0.411** | **< .001** |
| | | **Sponge** | **0.400** | **0.085** | **4.711** | **0.370** | **< .001** |
| | Clay | Sponge | -0.044 | 0.110 | -0.403 | -0.041 | 0.687 |

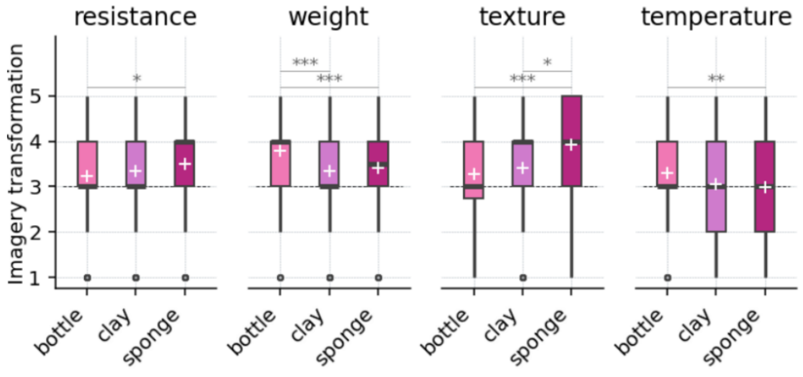


*Supplementary Figure 3 – OBJECT x PROPERTY interaction effect for active tactile imagery* ***transformation****. Conditional comparisons on factor PROPERTY. Scores range from 1 to 5, with higher scores denoting higher imagery transformation. Boxes represent the interquartile ranges (IQR (from the 25th (Q1) to the 75th (Q3) percentile); whiskers show the minimum within Q1 -1.5 times the IQR and maximum within Q3 + 1.5 times the IQR. The black lines show the median, and the white + the mean per condition. The dashed line in eachdenotes the scale midpoint (3).* ∘ *= outlier; *significant difference at* $p_{holm} \leq .05$*; **significant difference at* $p_{holm} \leq .01$*; ***significant difference at* $p_{holm} \leq .001$*.*

### Passive tactile imagery

#### ***Transformation*** *- additional pots-hoc testing*

##### *BODY SITE x SENSATION*

As we were also interested in differences in imagery ability between body sites, we additionally performed conditional comparisons on factor SENSATION ( Suppl.Table. 6). Interestingly, there were no significant differences between body sites for transformation of vibration. Transformation of pinching was significantly higher for the lip than all other body sites. For stroking however, transformation of the lip was only significantly higher than the back. Moreover, transformation of stroking was significantly higher for the foot than the back and the shin.

| | | Transformation | | | Pleasantness | | |
|---|---|---|---|---|---|---|---|
| | | t(135) | d | $p_{holm}$ | t(45) | d | $p_{holm}$ |
| **stroking** | | | | | | | |
| lip | foot | -7.83e$^{-15}$ | -4.44e$^{-16}$ | 1.000 | 0.15 | 0.03 | 1.000 |
| | shin | 2.07 | 0.21 | .122 | 1.55 | 0.25 | .643 |
| | back | **2.68** | **0.25** | **.038** | -1.14e$^{-15}$ | -2.22e$^{-16}$ | 1.000 |
| foot | shin | **2.71** | **0.21** | **.038** | 1.35 | 0.22 | .741 |
| | back | **2.79** | **0.25** | **.036** | -0.17 | -0.03 | 1.000 |
| shin | back | 0.44 | 0.04 | .000 | -1.91 | -0.25 | .376 |
| **pinching** | | | | | | | |
| lip | foot | **2.97** | **0.25** | **.014** | 0.18 | 0.03 | 1.000 |
| | shin | **4.73** | **0.33** | **< .001** | 0.04 | 0.01 | 1.000 |
| | back | **4.84** | **0.39** | **< .001** | -0.96 | -0.11 | 1.000 |
| foot | shin | 0.85 | 0.08 | .796 | -0.23 | -0.03 | 1.000 |
| | back | 1.40 | 0.13 | .493 | -1.03 | -0.14 | 1.000 |
| shin | back | 0.66 | 0.06 | .796 | -0.96 | -0.12 | 1.000 |
| **vibration** | | | | | | | |
| lip | foot | -0.69 | -0.07 | 1.000 | -2.02 | -0.43 | .135 |
| | shin | 1.41 | 0.14 | .639 | -0.32 | -0.05 | .753 |
| | back | -0.30 | -0.03 | 1.000 | **-4.16** | **-0.90** | **< .001** |
| foot | shin | 2.44 | 0.21 | .095 | 2.06 | 0.38 | .135 |
| | back | 0.41 | 0.04 | 1.000 | -2.58 | -0.47 | .053 |
| shin | back | -1.94 | -0.17 | .270 | **-4.11** | **-0.85** | **< .001** |

*Supplementary Table 6 Post-hoc pairwise comparisons for BODY SITE x SENSATION interaction effects for passive tactile imagery* ***transformation*** *and* ***pleasantness****. Conditional on factor SENSATION. d = Cohen's d; $p_{holm}$ = Holm-Bonferroni corrected p-value adjusted for comparing a family of 6. Significant differences are printed in bold.*

#### ***Pleasantness*** *- Additional post-hocs*

##### *BODY SITE*

*Supplementary Table 7 - Post Hoc comparisons for significant main effect of BODY SITE for imagery* ***pleasantness****. Significant differences are printed in bold. $p_{holm}$ = Holm-Bonferroni corrected p-value adjusted for comparing a family of 6; SE = standard error.*

| | | mean difference | SE | t(45) | Cohen's d | $p_{holm}$ |
|---|---|---|---|---|---|---|
| lip | foot | -0.500 | 0.540 | -0.926 | -0.123 | 0.719 |
| | shin | 0.275 | 0.341 | 0.809 | 0.068 | 0.719 |
| | back | -1.377 | 0.562 | -2.450 | -0.338 | 0.091 |
| foot | shin | 0.775 | 0.434 | 1.786 | 0.190 | 0.242 |
| | back | -0.877 | 0.445 | -1.972 | -0.215 | 0.219 |
| **shin** | **back** | **-1.652** | **0.418** | **-3.952** | **-0.405** | **0.002** |

Supplementary materials to *Capturing the imagination: mapping imagery ability across our multidimensional sense of touch* – Lustenhouwer & Dijkerman

#### *BODY SITE x SENSATION*

We additionally performed conditional comparisons based on factor SENSATION to allow comparison of body sites for each of the sensations (Suppl.Table.6). Interestingly, pleasantness did not differ between body sites for stroking and pinching. For vibration however, pleasantness was significantly higher for the back than the lip and the shin and showed a trend in the same direction for the back vs. the foot.

#### *Non-significant correlations between imagery pleasantness and ability per passive sensation*

*Supplementary Table 8 – non-significant correlations between imagery ability (imagery processing components vividness, maintenance and transformation) and pleasantness for pinching and vibration.* $p_{holm}$ *= Holm-Bonferroni corrected p-value adjusted for comparing a family of 3.*

| IMAGERY PROCESSING COMPONENTS | PINCHING | VIBRATION |
|---|---|---|
| VIVIDNESS | $\rho(44) = -.12$, $p_{holm} = .836$ | $\rho(44) = .20$, $p_{holm} = .243$ |
| MAINTENANCE | $r(44) = -.07$, $p_{holm} = .836$ | $\rho(44) = .23$, $p_{holm} = .243$ |
| TRANSFORMATION | $\rho(44) = -.20$, $p_{holm} = .548$ | $\rho(44) = .34$, $p_{holm} = .064$ |
